# Scalable Bayesian GPFA with automatic relevance determination and discrete noise models

**DOI:** 10.1101/2021.06.03.446788

**Authors:** Kristopher T. Jensen, Ta-Chu Kao, Jasmine T. Stone, Guillaume Hennequin

## Abstract

Latent variable models are ubiquitous in the exploratory analysis of neural population recordings, where they allow researchers to summarize the activity of large populations of neurons in lower dimensional ‘latent’ spaces. Existing methods can generally be categorized into (i) Bayesian methods that facilitate flexible incorporation of prior knowledge and uncertainty estimation, but which typically do not scale to large datasets; and (ii) highly parameterized methods without explicit priors that scale better but often struggle in the low-data regime. Here, we bridge this gap by developing a fully Bayesian yet scalable version of Gaussian process factor analysis (bGPFA), which models neural data as arising from a set of inferred latent processes with a prior that encourages smoothness over time. Additionally, bGPFA uses automatic relevance determination to infer the dimensionality of neural activity directly from the training data during optimization. To enable the analysis of continuous recordings without trial structure, we introduce a novel variational inference strategy that scales near-linearly in time and also allows for non-Gaussian noise models appropriate for electrophysiological recordings. We apply bGPFA to continuous recordings spanning 30 minutes with over 14 million data points from primate motor and somatosensory cortices during a self-paced reaching task. We show that neural activity progresses from an initial state at target onset to a reach-specific preparatory state well before movement onset. The distance between these initial and preparatory latent states is predictive of reaction times across reaches, suggesting that such preparatory dynamics have behavioral relevance despite the lack of externally imposed delay periods. Additionally, bGPFA discovers latent processes that evolve over slow timescales on the order of several seconds and contain complementary information about reaction time. These timescales are longer than those revealed by methods which focus on individual movement epochs and may reflect fluctuations in e.g. task engagement.

## 1 Introduction

The adult human brain contains upwards of 100 billion neurons (Azevedo et al., 2009). Yet many of our day-to-day behaviors such as navigation, motor control, and decision making can be described in much lower dimensional spaces. Accordingly, recent studies across a range of cognitive and motor tasks have shown that neural population activity can often be accurately summarised by the dynamics of a “latent state” evolving in a low-dimensional space (Churchland et al., 2012; Pandarinath et al., 2018; Chaudhuri et al., 2019; Minxha et al., 2020; Ecker et al., 2014). Inferring and investigating these latent processes can therefore help us understand the underlying representations and computations implemented by the brain (Humphries, 2020). To this end, numerous latent variable models have been developed and used to analyze the activity of populations of simultaneously recorded neurons. These models range from simple linear projections such as PCA to sophisticated non-linear and temporally correlated models (Jensen et al., 2020; Pandarinath et al., 2018; Gao et al., 2016; Cunningham and Byron, 2014; Schimel et al., 2021).

A popular latent variable model for neural data analysis is Gaussian process factor analysis (GPFA), which has yielded insights into neural computations ranging from time tracking to movement preparation and execution (Afshar et al., 2011; Sohn et al., 2019; Sauerbrei et al., 2020; Rutten et al., 2020). However, fitting GPFA comes with a computational complexity of 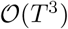 and a memory footprint 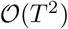 for *T* time bins. This prohibits the application of GPFA to time series longer than a few hundred time bins without artificially chunking such data into “pseudo-trials” and treating these as independent samples. Additionally, canonical GPFA assumes a Gaussian noise model while recent work has suggested that non-Gaussian models often perform better on neural data (Duncker and Sahani, 2018; Keeley et al., 2020b; Zhao and Park, 2017). Here, we address these challenges by formulating a scalable and fully Bayesian version of GPFA (bGPFA; Figure 1) with a computational complexity of 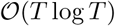 and a memory cost of 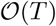. To do this, we introduce an efficiently parameterized variational inference strategy that ensures scalability to long recordings while also supporting non-Gaussian noise models. Additionally, the Bayesian formulation provides a framework for principled model selection based on approximate marginal likelihoods (Titsias and Lawrence, 2010). This allows us to perform automatic relevance determination and thus fit a single model without prior assumptions about the underlying dimensionality, which is instead inferred from the data itself (Neal, 2012; Bishop, 1999).

**Figure 1:**
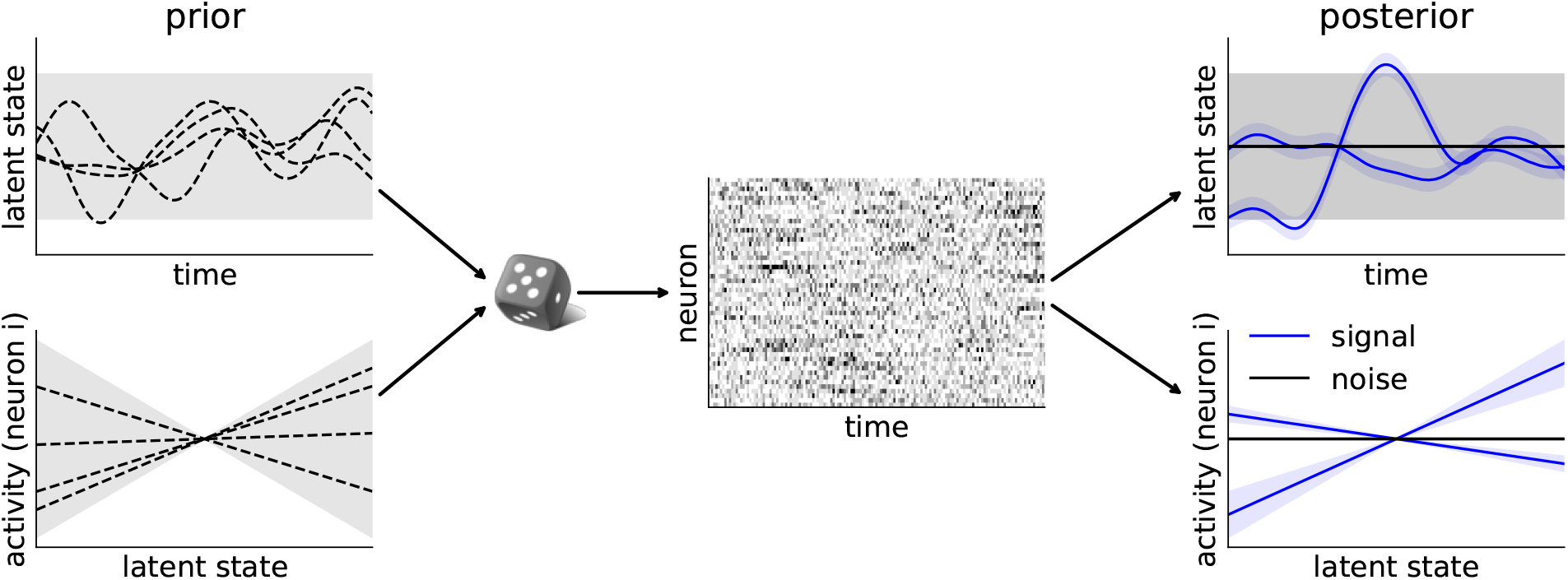
Bayesian GPFA schematic. Bayesian GPFA places a Gaussian Process prior over the latent states in each dimension as a function of time ***t*** (*p*(***X***|***t***); top left) as well as a linear prior over neural activity as a function of each latent dimension (*p*(***F***|***X***); bottom left). Together with a stochastic noise process *p*(***Y***|***F***), which can be discrete for electrophysiological recordings, this forms a generative model that gives rise to observations ***Y*** (middle). From the data and priors, bGPFA infers posterior latent states for each latent dimension (*p*(***X***|***Y***); top right) as well as a posterior predictive observation model for each neuron (*p*(***Y***_*test*_|***X***_*test*_, ***Y***); bottom right). When combined with automatic relevance determination, the model learns to automatically discard superfluous latent dimensions by maximizing the log marginal likelihood of the data (right, black vs. blue).

We validate our method on synthetic and biological data, where bGPFA exhibits superior performance to GPFA and Poisson GPFA with increased scalability and without requiring cross-validation to select the latent dimensionality. We then apply bGPFA to longitudinal, multi-area recordings from primary motor (M1) and sensory (S1) areas during a monkey self-paced reaching task spanning 30 minutes. bGPFA readily scales to such datasets, and the inferred latent trajectories improve decoding of kinematic variables compared to the raw data. This decoding improves further when taking into account the temporal offset between motor planning encoded by M1 and feedback encoded by S1. We also show that the latent trajectories for M1 converge to consistent regions of state space for a given reach direction at the onset of each individual reach. Importantly, the distance in latent space to this preparatory state from the state at target onset is predictive of reaction times across reaches, similar to previous results in a task that includes an explicit ‘motor preparation epoch’ where the subject is not allowed to move (Afshar et al., 2011). This illustrates the functional relevance of such preparatory activity and suggests that motor preparation takes place even when the task lacks well-defined trial structure and externally imposed delay periods, consistent with findings by Lara et al. (2018) and Zimnik and Churchland (2021). Finally, we analyze the task relevance of slow latent processes identified by bGPFA which evolve on timescales of seconds; longer than the millisecond timescales that can be resolved by methods designed for trial-structured data. We find that some of these slow processes are also predictive of reaction time across reaches, and we hypothesize that they reflect task engagement which varies over the course of several reaches.

## 2 Method

In the following, we use the notation ***A*** to refer to the matrix with elements *a_ij_*. We use ***a***_*k*_ to refer to the k^th^ row *or* column of ***A*** with an index running from 1 to *K*, represented as a column vector.

### 2.1 Generative model

Latent variable models for neural recordings typically model the neural activity 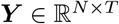 of *N* neurons at times 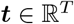 as arising from shared fluctuations in *D* latent variables 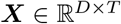. Specifically, the probability of a given recording can be written as

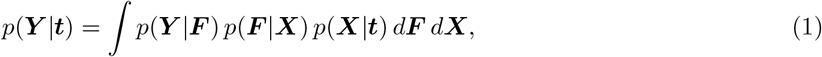

where 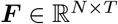 are intermediate, neuron-specific variables that can often be thought of as firing rates or a similar notion of noise-free activity. For example, GPFA (Yu et al., 2009) specifies

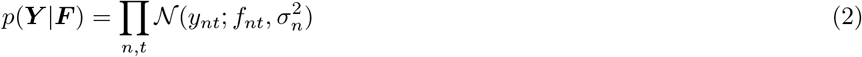

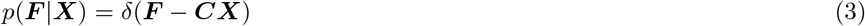

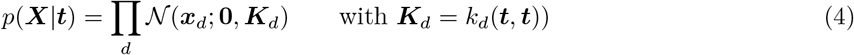

That is, the prior over the *d*^th^ latent function *x_d_*(*t*) is a Gaussian process (Rasmussen and Williams, 1996) with covariance function *k_d_*(·, ·) (usually a radial basis function), and the observation model *p*(***Y***|***X***) is given by a parametric linear transformation with independent Gaussian noise.

In this work, we additionally introduce a prior distribution over the mixing matrix 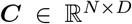 with hyperparameters specific to each latent dimension. This allows us to *learn* an appropriate latent dimensionality for a given dataset using automatic relevance determination (ARD) similar to previous work in Bayesian PCA (Appendix I; Bishop, 1999) rather than relying on cross-validation or ad-hoc thresholds of variance explained. Unlike in standard GPFA, the log marginal likelihood (Equation 1) becomes intractable with this prior. We therefore develop a novel variational inference strategy (Wainwright and Jordan, 2008) which also (i) provides a scalable implementation appropriate for long continuous neural recordings, and (ii) extends the model to general non-Gaussian likelihoods better suited for discrete spike counts.

In this new framework, which we call Bayesian GPFA (bGPFA), we use a Gaussian prior over ***C*** of the form 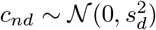, where *s_d_* is a scale parameter associated with latent dimension *d*. Integrating ***C*** out in Equation 3 then yields the following observation model:

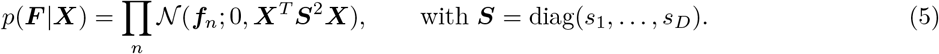

Moreover, we use a general noise model 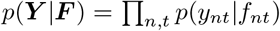 where *p*(*y_nt_*|*f_nt_*) is any distribution for which we can evaluate its density.

### 2.2 Variational inference and learning

To train the model and infer both ***X*** and ***F*** from the data ***Y***, we use a nested variational approach. It is intractable to compute log *p*(***Y***|***t***) (Equation 1) analytically for bGPFA, and we therefore introduce a lower bound on log *p*(***Y***|***t***) at the outer level and another one on log *p*(***Y***|***X***) at the inner level. These lower bounds are constructed from approximations to the posterior distributions over latents (***X***) and noise-free activity (***F***) respectively.

#### Distribution over latents

At the outer level, we introduce a variational distribution *q*(***X***) over latents and construct an evidence lower bound (ELBO; Wainwright and Jordan, 2008) on the log marginal likelihood of Equation 1:

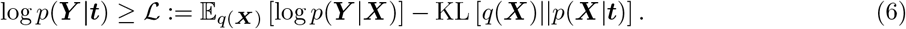

Conveniently, maximizing this lower bound is equivalent to minimizing KL [*q*(***X***)‖*p*(***X***|***Y***)] and thus also yields an approximation to the posterior over latents in the form of *q*(***X***). We estimate the first term of the ELBO using Monte Carlo samples from *q*(***X***) and compute the KL term analytically.

Here, we use a so-called whitened parameterization of *q*(***X***) (Hensman et al., 2015b) that is both expressive and scalable to large datasets:

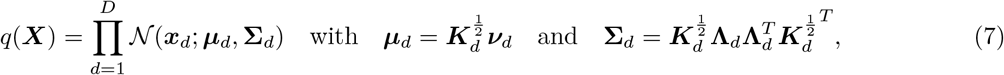

where 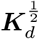 is any square root of the prior covariance matrix ***K**_d_*, and 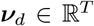 is a vector of variational parameters to be optimized. 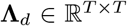 is a positive semi-definite variational matrix whose structure is chosen carefully so that its squared Frobenius norm, log determinant, and matrix-vector products can all be computed efficiently, which facilitates the evaluation of Equations 8 and 9. This whitened parameterization has several advantages. First, it does not place probability mass where the prior itself does not. In addition to stabilizing learning (Murray and Adams, 2010), this also guarantees that the posterior is temporally smooth for a smooth prior. Second, the KL term in Equation 6 simplifies to

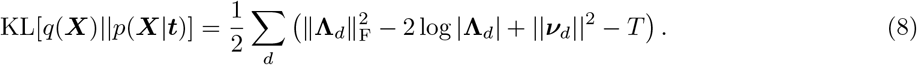

Third, *q*(***X***) can be sampled efficiently via a differentiable transform (i.e. the reparameterization trick) provided that fast differentiable 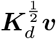 and **Λ**_*d*_***v*** products are available for any vector ***v***:

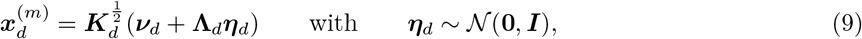

where 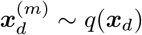. This is important to form a Monte Carlo estimate of 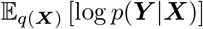.

To avoid the challenging computation of 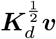 for general ***K**_d_* (Allen et al., 2000), we directly parameterize 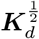, the positive definite square root of ***K***, which implicitly defines the prior covariance function *k_d_*(·, ·). In this work we use an RBF kernel for ***K**_d_* and give the expression for 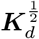 in Appendix F. Additionally, we use Toeplitz acceleration methods to compute 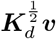 products in 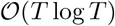 time and with 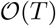 memory cost (Wilson et al., 2015; Rutten et al., 2020).

We implement and compare different choices of **Λ***_d_* in Appendix F. For the experiments in this work, we use the parameterization **Λ***_d_* = **Ψ***_d_**C**_d_*, where **Ψ***_d_* is diagonal with positive entries and ***C**_d_* is circulant, symmetric, and positive definite. This parameterization enables cheap computation of KL divergences and matrix-vector products while maintaining sufficient expressiveness (Appendix F). All results are qualitatively similar when instead using a simple diagonal parameterization **Λ***_d_* = **Ψ***_d_*.

#### Distribution over neural activity

Evaluating 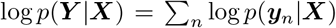 for each sample drawn from *q*(***X***) is intractable for general noise models. Thus, we further lower-bound the ELBO of Equation 6 by introducing an approximation *q*(***f***_*n*_|***X***) to the posterior *p*(***f**_n_*|***y**_n_*, ***X***):

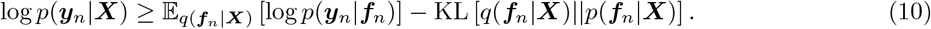

We repeat the whitened variational strategy described at the outer level by writing

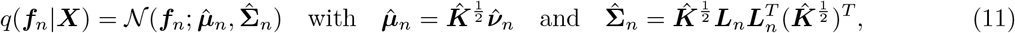

where 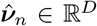 is a neuron-specific vector of variational parameters to be optimized along with a lower-triangular matrix 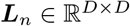; and 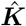 denotes the covariance matrix of *p*(***f*** |***X***), whose square root 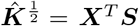 follows from Equation 5. The low-rank structure of 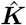 enables cheap matrix-vector products and KL divergences:

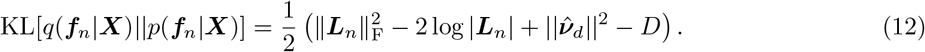

Note that the KL divergence does not depend on ***X*** in this whitened parameterization (Appendix H). Moreover, *q*(***f***_*n*_|***X***) in Equation 11 has the form of the exact posterior when the noise model is Gaussian (Appendix G), and it is equivalent to a stochastic variational inducing point approximation (Hensman et al., 2015a) for general noise models (Appendix H).

Finally, we need to compute the first term in Equation 10:

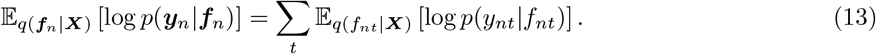

Each term in this sum is simply a 1-dimensional Gaussian expectation which can be computed analytically in the case of Gaussian or Poisson noise (with an exponential link function), and otherwise approximated efficiently using Gauss-Hermite quadrature (Appendix K; Hensman et al., 2015a).

### 2.3 Summary of the algorithm

Putting Section 2.1 and Section 2.2 together, optimization proceeds at each iteration by drawing *M* Monte Carlo samples 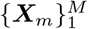 from *q*(***X***) and estimating the overall ELBO as:

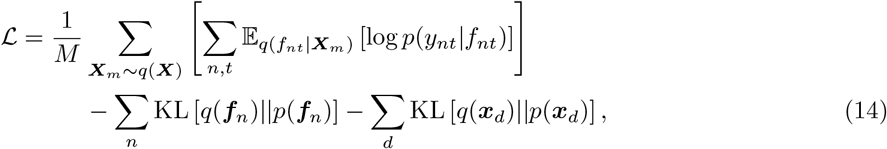

where the expectation over *q*(*f_nt_*|***X***) is evaluated analytically or using Gauss-Hermite quadrature depending on the noise model (Appendix K). We maximize 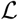 using stochastic gradient ascent with Adam (Kingma and Ba, 2015). This has a total computational time complexity of 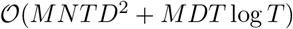 and memory complexity of 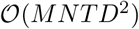, where *N* is the number of neurons, *T* the number of time points, and *D* the latent dimensionality. For large datasets such as the monkey reaching data in Section 3.2, we compute gradients using mini-batches across time to mitigate the memory cost. That is, gradients for the sum over *t* in Equation 14 are computed in multiple passes. The algorithm is described in pseudocode with further implementation and computational details in Appendix L. The model learned by bGPFA can subsequently be used for predictions on held-out data by conditioning on partial observations as used for cross-validation in Section 3.1 and discussed in Appendix M. Latent dimensions that have been ‘discarded’ by automatic relevance determination will automatically have negligible contributions to the resulting posterior predictive distribution since the prior scale parameters *s_d_* are approximately zero for these dimensions (see Appendix I for details).

## 3 Experiments and results

### 3.1 Synthetic data

We first generated an example dataset from the GPFA generative model (Equations 2–4) with a true latent dimensionality of 3. We proceeded to fit both factor analysis (FA), GPFA, and bGPFA with different latent dimensionalities *D* ∈ [1, 10]. Here, we fitted bGPFA without automatic relevance determination such that *s_d_* = *s* ∀*d*. As expected, the marginal likelihoods increased monotonically with *D* for both FA and GPFA (Figure 2a; Appendix I). In contrast, the bGPFA ELBO reached its optimum value at the true latent dimensionality *D*^⋆^ = 3. This is a manifestation of “Occam’s razor”, whereby fully Bayesian approaches favor the simplest model that adequately explains the data ***Y*** (MacKay, 2003). When instead considering the cross-validated predictive performance of each method, performance deteriorated rapidly for *D* > 3 for FA and GPFA, while Bayesian GPFA was more robust to overfitting (Figure 2b). Notably, the introduction of ARD parameters {*s_d_*} in bGPFA allowed us to fit a single model with large *D* = 10. This recovered the maximum ELBO of bGPFA without ARD and the minimum test error across GPFA and bGPFA without ARD (Figure 2a and b, black) without *a priori* assumptions about the latent dimensionality or the need to perform extensive cross-validation. Consistent with the ground truth generative process, only 3 of the scale parameters *s_d_* remained well above zero after training (Figure 2b, inset). Similar to this illustrative example with Gaussian data, bGPFA with ARD and Poisson noise also exceeded the optimal performance of Poisson GPFA when applied to both synthetic and experimental spike count data (Appendix E).

**Figure 2:**
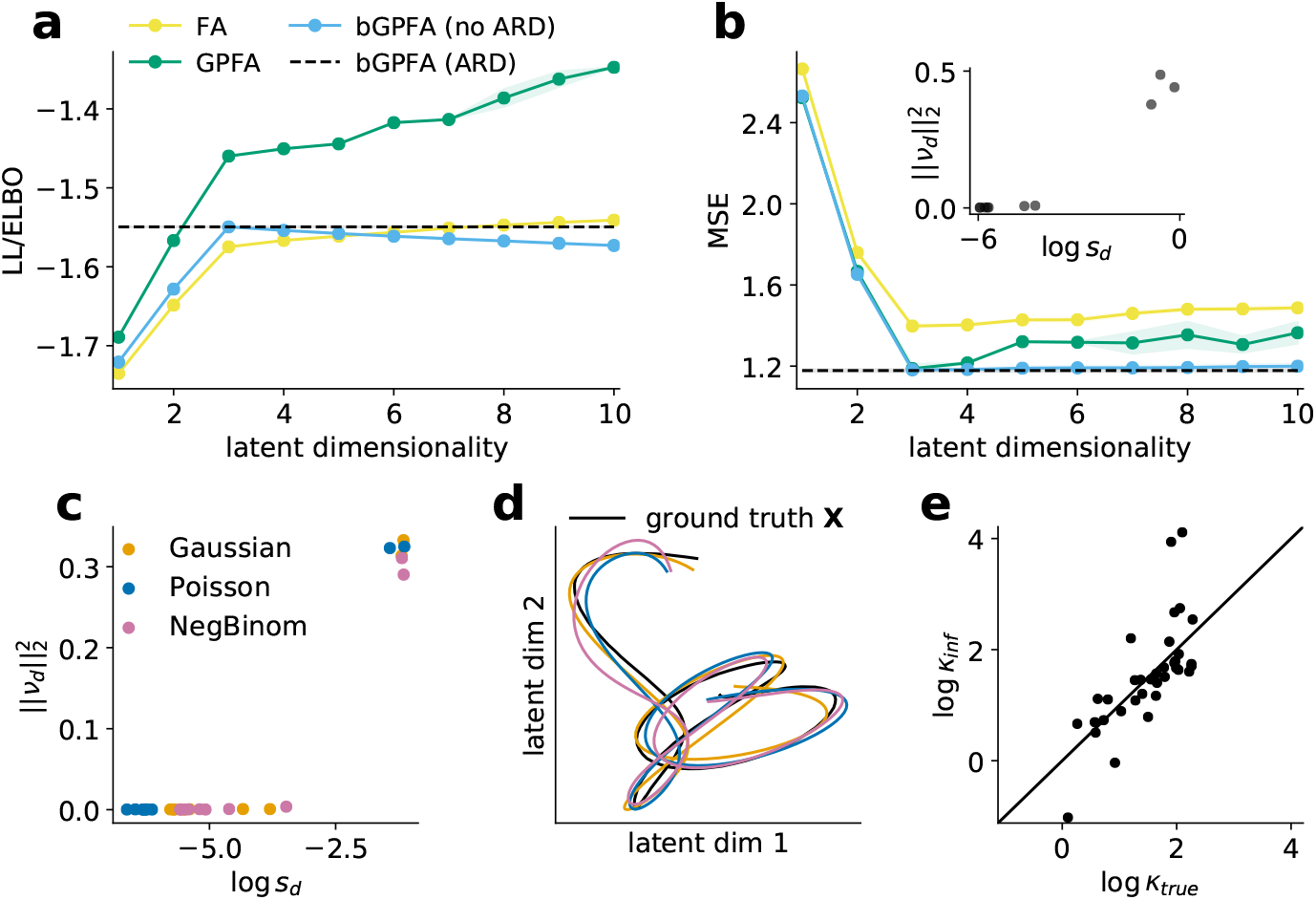
Bayesian GPFA applied to synthetic data. **(a)** Log likelihoods of factor analysis (yellow) & GPFA (green) and ELBO of Bayesian GPFA without ARD (blue) fitted to synthetic data with a ground truth dimensionality of three for different model dimensionalities. bGPFA with ARD recovered a three-dimensional latent space as well as the optimum ELBO of bGPFA without ARD (black dashed line). **(b)** Cross-validated prediction errors for the models in (a) (Appendix M). bGPFA with ARD recovered the performance of the optimal GPFA and bGPFA models without requiring a search over latent dimensionalities. Inspection of the learned prior scales {*s_d_*} and posterior mean parameters 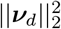 (inset) indicates that ARD retained only *D*^⋆^ = 3 informative dimensions (top right) and discarded the other 7 dimensions (bottom left). Shadings in (a) and (b) indicate ±2 stdev. across 10 model fits. **(c)** Learned parameters of bGPFA with ARD and either Gaussian, Poisson or negative binomial noise models fitted to two-dimensional synthetic datasets with observations drawn from the corresponding noise models (Appendix K). The parameters clustered into two groups of informative (top right) and non-informative (bottom left) dimensions (Appendix J). **(d)** Latent trajectory in the space of the two most informative dimensions (c.f. (c)) for each model with the ground truth shown in black. **(e)** The overdispersion parameter *κ_n_* for each neuron learned in the negative binomial model, plotted against the ground truth (Appendix K). Solid line indicates *y* = *x*; note that *κ_n_* → ∞ corresponds to a Poisson noise model.

We then proceeded to apply bGPFA (*D* = 10) to an example dataset drawn using Equations 4 and 5 with a ground truth dimensionality *D*^⋆^ = 2, and either Gaussian, Poisson, or negative binomial noise. For all three datasets, the learned parameters clustered into a group of two latent dimensions with high information content (Appendix J) and a group of eight uninformative dimensions, consistent with the generative process (Figure 2c). In each case, we extracted the inferred latent trajectories corresponding to the informative dimensions and found that they recapitulated the ground truth up to a linear transformation (Figure 2d). Fitting flexible noise models such as the negative binomial model is important because neural firing patterns are known to be overdispersed in many contexts (Tomko and Crapper, 1974; Fenton and Muller, 1998; Azouz and Gray, 1999). However, it is often unclear how much of that overdispersion should be attributed to common fluctuations in hidden latent variables (***X*** in our model) compared to private noise processes in single neurons (Low et al., 2018). In our synthetic data with negative binomial noise, we could accurately recover the single-neuron overdispersion parameters (Figure 2e; Appendix K), suggesting that such unsupervised models have the capacity to resolve overdispersion due to private and shared processes.

In summary, bGPFA provides a flexible method for inferring both latent dimensionalities, latent trajectories, and heterogeneous single-neuron parameters in an unsupervised manner. In the next section, we show that the scalability of the model and its interpretable parameters also facilitate the analysis of large neural population recordings.

### 3.2 Primate recordings

In this section, we apply bGPFA to biological data recorded from a rhesus macaque during a self-paced reaching task with continuous recordings spanning 30 minutes (O’Doherty et al., 2017; Makin et al., 2018; Figure 3a). The continuous nature of these recordings as one long trial makes it a challenging dataset for existing analysis methods that explicitly require the availability of many trials per experimental condition (Pandarinath et al., 2018), and poses computational challenges to Gaussian process-based methods that cannot handle long time series (Yu et al., 2009). While the ad-hoc division of continuous recordings into surrogate trials can still enable the use of these methods (Keshtkaran et al., 2021), here we show that our formulation of bGPFA readily applies to long continuous recordings. We fitted bGPFA with a negative binomial noise model to recordings from both primary motor cortex (M1) and primary somatosensory cortex (S1). For all analyses, we used a single recording session (`indy_20160426`, as in Keshtkaran et al., 2021), excluded neurons with overall firing rates below 2 Hz, and binned data at 25 ms resolution. This resulted in a data array 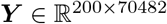 (130 M1 neurons and 70 S1 neurons).

**Figure 3:**
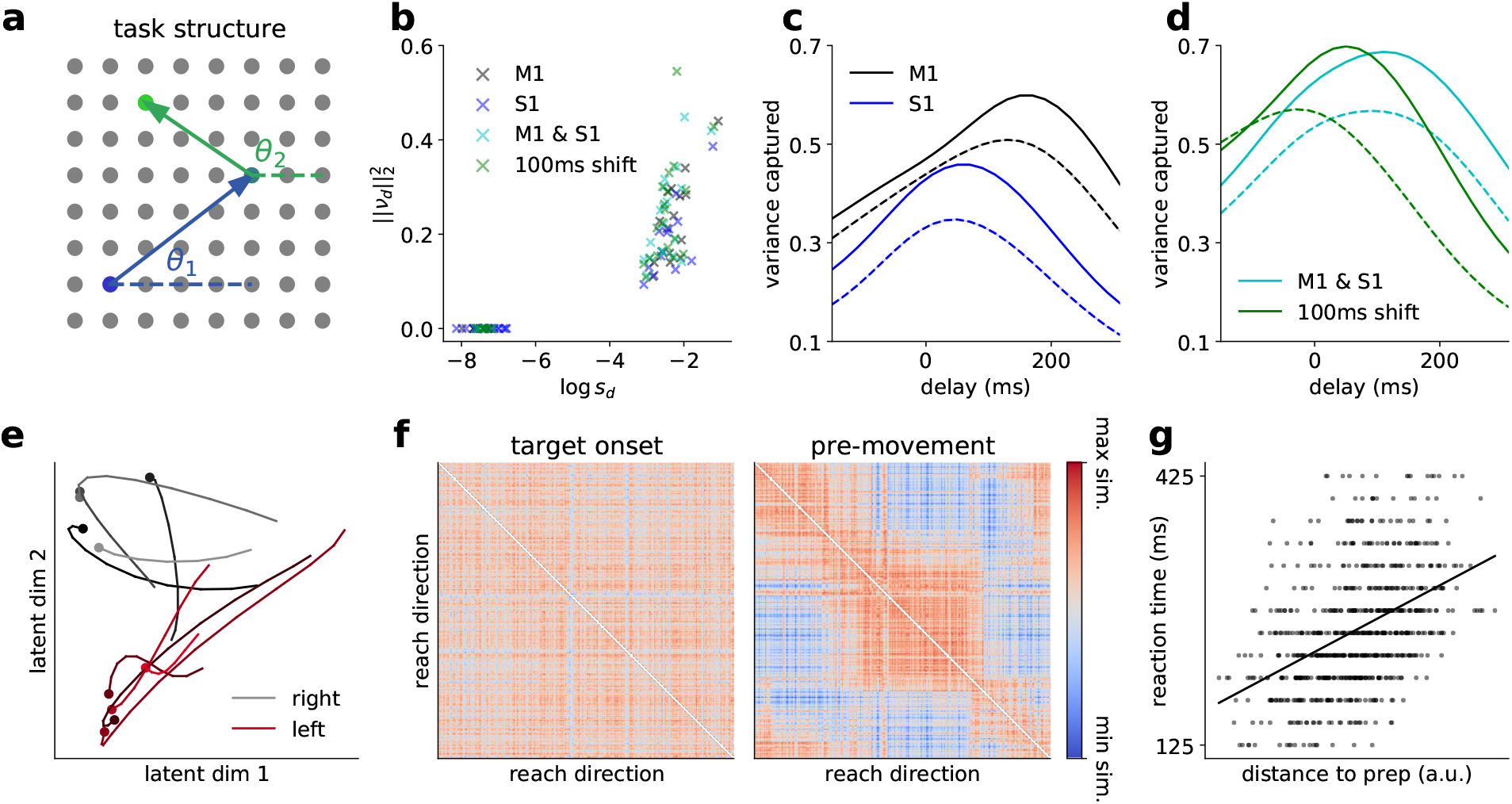
Bayesian GPFA applied to primate data. **(a)** Schematic illustration of the self-paced reaching task. When a target on a 17×8 grid is reached (arrows; 8×8 shown for clarity), a new target lights up on the screen (colours), selected at random from the remaining targets. In several analyses, we classify movements according to reach angle measured relative to horizontal (*θ*_1_, *θ*_2_). **(b)** Learned mean and scale parameters for the bGPFA models. Small prior scales *s_d_* and posterior mean parameters 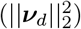 indicate uninformative dimensions (Appendix J). **(c)** We applied bGPFA to monkey M1 and S1 data during the task and trained a linear model to decode kinematics from firing rates predicted from the inferred latent trajectories with different delays between latent states and kinematics. Neural activity was most predictive of future behavior in M1 (black) and current behavior in S1 (blue). Dashed lines indicate decoding from the raw data convolved with a Gaussian filter. **(d)** Decoding from bGPFA applied to the combined M1 and S1 data (cyan). Performance improved further when decoding from latent trajectories inferred from data where M1 activity was shifted by 100 ms relative to S1 activity (green). **(e)** Example trajectories in the two most informative latent dimensions for five rightward reaches (grey) and five leftward reaches (red). Trajectories are plotted from the appearance of the stimulus until movement onset (circles). During ‘movement preparation’, the latent trajectories move towards a consistent region of latent state space for each reach direction. **(f)** Similarity matrix of the latent state at stimulus onset showing no obvious structure (left) and 75 ms prior to movement onset showing modulation by reach direction (right). **(g)** Reaction time plotted against Euclidean distance between the latent state at target onset and the mean preparatory state for the corresponding reach direction (*ρ* = 0.45).

We first fitted bGPFA independently to the M1 and S1 sub-populations with *D* = 25 latent dimensions. In this case, ARD retained 16 (M1) and 12 (S1) dimensions (Figure 3b). We then proceeded to train a linear decoder to predict hand kinematics in the form of *x* and *y* hand velocities from either the inferred firing rates or the raw data convolved with a 50 ms Gaussian kernel (Keshtkaran et al., 2021; Appendix M). We found that the model learned by bGPFA predicted kinematics better than the convolved spike trains, suggesting that (i) the latent space accurately captures kinematic representations, and (ii) the denoising and data-sharing across time in bGPFA aids decodability beyond simple smoothing of neural activity. Interestingly, by repeating this decoding analysis with an artificially imposed delay between neural activity and decoded behavior, we found that neurons in S1 predominantly encoded current behavior while neurons in M1 encoded a motor plan that predicted kinematics 100-150 ms into the future (Figure 3b). This is consistent with the motor neuroscience literature suggesting that M1 functions as a dynamical system driving behavior via downstream effectors (Churchland et al., 2012).

We then fitted bGPFA to the entire dataset including both M1 and S1 neurons. In this case, bGPFA retained 19 dimensions (Appendix D), and kinematic predictions improved over individual M1- and S1-based predictions (Figure 3b). In this analysis, the decoding performance as a function of delay between neural activity and behavior exhibited a broader peak than for the single-region decoding. We hypothesized that this broad peak reflects the fact that these neural populations encode both *current* behavior in S1 as well as *future* behavior in M1 (Figure 3c). Indeed, when we took this offset into account by shifting all M1 spike times by +100 ms and retraining the model, decoding performance increased from 68.56% ± 0.09 to 69.81% ± 0.06 (mean ± sem variance explained across ten model fits; Appendix M). Additionally, the shifted data exhibited a narrower decoding peak attained for near-zero delay between kinematics and latent trajectories (Figure 3d). Consistent with the improved kinematic decoding, we also found that shifting the M1 spikes by 100 ms increased the ELBO per neuron (−34,637.0 ± 0.7 to −34,631.1 ± 0.6) and decreased the dimensionality of the data (Appendix D; Recanatesi et al., 2019). These results suggest that M1 and S1 contain both overlapping but also non-redundant information, and that the most parsimonious description of the neural data is recovered by taking into account the different biological properties of M1 and S1.

We next wondered if bGPFA could be used to reveal putative motor preparation processes, which is non-trivial due to the lack of trial structure and well-defined preparatory epochs. We partitioned the data post-hoc into individual ‘reaches’, each consisting of a period of time where the target location remained constant. For these analyses, we only considered ‘successful’ reaches, where the monkey eventually moved to the target location, and we defined movement onset as the first time during a reach where the cursor speed exceeded a low threshold (Appendix A). We began by visualizing the latent processes inferred by bGPFA as they unfolded prior to movement onset in each reach epoch. For visualization purposes, we ranked the latent dimensions based on their learned prior scales (a measure of variance explained; Appendix J) and selected the first two. Prior to movement onset, the latent trajectories tended to progress from their initial location at target onset towards reach-specific regions of state space (see example trials in Figure 3e for leftward and rightward reaches). To quantify this phenomenon, we computed pairwise similarities between latent states across all 762 reaches, during (i) stimulus onset and (ii) 75 ms before movement onset (chosen such that it is well before any detectable movement; Appendix A). We defined similarity as the negative Euclidean distance between latent states and restricted the analysis to ‘fast’ latent dimensions with timescales smaller than 200 ms to study this putatively fast process. When plotted as a function of reach direction, the latent similarities at target onset showed little discernable structure (Figure 3f, left). In contrast, the pairwise similarities became strongly structured 75 ms before movement onset where neighboring reach directions were associated with similar preparatory latent states (Figure 3f, right). Similar albeit noisier results were found when using factor analysis or GPFA instead of bGPFA (Appendix A). These findings are consistent with previous reports of monkey M1 partitioning preparatory and movement-related activity into distinct subspaces (Elsayed et al., 2016; Lara et al., 2018), as well as with the analogous finding that a ‘relative target’ subspace is active before a ‘movement subspace’ in previous analyses of this particular dataset (Keshtkaran et al., 2021).

Previous work on delayed reaches has shown that monkeys start reaching earlier when the neural state attained at the time of the go cue – which marks the end of a delay period with a known reach direction – is close to an “optimal subspace” (Afshar et al., 2011; Kao et al., 2021). We wondered if a similar effect takes place during continuous, self-initiated reaching in the absence of explicit delay periods. Based on Figure 3e, we hypothesized that the monkey should start moving earlier if, at the time the next target is presented, its latent state is already close to the mean preparatory state for the required next movement direction. To test this, we extracted the mean preparatory state 75 ms prior to movement onset (as above) for each reach direction in the dataset. We found that the distance between the latent state at target onset and the corresponding mean preparatory state was strongly predictive of reaction time (Figure 3g, Pearson *ρ* = 0.45, *p* =4 × 10^−36^). Such a correlation was also weakly present with factor analysis (*ρ* = 0.21, *p* =1.1 × 10^−8^) but not detectable in the raw data (*ρ* = 0.002, *p* = 0.95). We also verified that the strong correlation found with bGPFA was not an artifact of the temporal correlations introduced by the prior (Appendix C). Taken together, our results suggest that motor preparation is an important part of reaching movements even in an unconstrained self-paced task. Additionally, we showed that bGPFA captures such behaviorally relevant latent dynamics better than simpler alternatives, and our scalable implementation enables its use on the large continuous reaching dataset analysed here.

### 3.3 Long-timescale latent processes

Some latent dimensions inferred by bGPFA also had long timescales on the order of 1.5 seconds, which is similar to the timescale of individual reaches (1-2 seconds; Appendix C). We hypothesized that these slow dynamics might reflect motivation or task engagement. Consistent with this hypothesis, we found that one of the slow latent processes (*τ* = 1.4 s) was strongly correlated with reaction time during successful reaches (Pearson *ρ* = 0.40, *p* = 3.4 × 10^−28^). Interestingly, the information contained about reaction time in this long timescale latent dimension was largely complementary to that encoded by the distance to preparatory states in the ‘fast’ dimensions (Appendix C), suggesting that motor preparation and task engagement are orthogonal processes both contributing to task performance.

The experimental recordings were also characterized by a period of approximately five minutes towards the end of the recording session during which the monkey did not participate actively in the task and the cursor velocity was near-constant at zero (Figure 4a). When analysing neural activity across the periods with and without task participation, we found that neural dynamics moved to a different subspace as the monkey stopped engaging with the task (Figure 4b). Importantly, we were able to simultaneously capture these context-dependent changes as well as movement-specific and preparatory dynamics (Section 3.2) by fitting a single model to the full 30 minute dataset. This suggests that bGPFA can capture behaviorally relevant dynamics within individual contexts even when trained on richer datasets with changing contexts.

**Figure 4:**
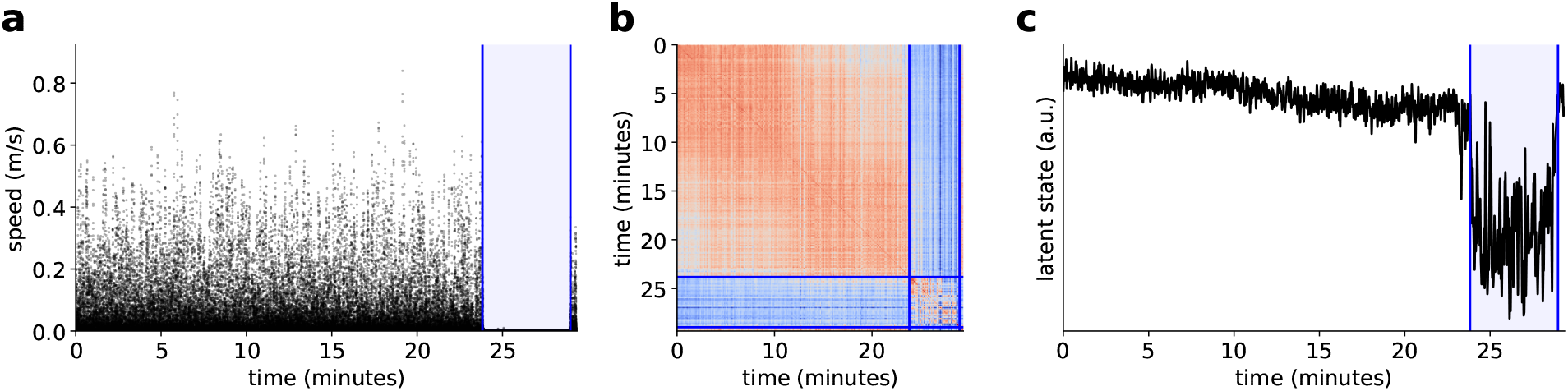
Analysis of a period without task participation. **(a)** Cursor speed over the course of the recording session. Blue horizontal lines indicate the last successful trial before and first successful trial after a period with no active task participation (blue shading). **(b)** Latent similarity matrix as a function of time during the task. The latent dynamics during task participation occur in a largely orthogonal subspace to the dynamics during the period with no active task participation. **(c)** Plot of latent state over time for a long-timescale latent dimension strongly correlated with reaction time (*τ* = 1.4 s).

Finally, we wondered how the neural activity patterns during periods with and without task participation were related to the long-timescale latent dimensions predictive of task engagement. Here we found that the slow latent process considered above also exhibited a prominent change to a different state as the monkey stopped participating in the task (Figure 4). This is consistent with our hypothesis that this latent process captures a feature related to task engagement which slowly deteriorated during the first 24 minutes of the task followed by a discrete switch to a state with no engagement in the task. During the period of active task participation, this latent dimension was also correlated with time within the session. Indeed, reach number and latent state were both predictive of reaction time, but with the latent trajectory exhibiting a slightly stronger correlation (Pearson *ρ* = 0.40 vs. *ρ* = 0.37). It is not surprising that task engagement decreases with time, and it is in this case difficult to tease apart how motivation and time are differentially represented in such latent processes. However, based on the strong and abrupt modulation by task participation, this latent dimension appears to represent an aspect of engagement with the task beyond the passing of time.

Taken together, we thus find that bGPFA is capable of capturing not only single-reach dynamics and preparatory activity but also complementary processes evolving over longer timescales, which would be difficult to identify with methods designed for the analysis of many shorter trials.

## 4 Discussion

### Related work

The generative model of bGPFA can be considered an extension of the canonical GPFA model proposed by Yu et al. (2009) to include a Gaussian prior over the loading matrix ***C*** (Section 2.1). In this view, bGPFA is to GPFA what Bayesian PCA is to PCA (Bishop, 1999); in particular, it facilitates automatic relevance determination to infer the dimensionality of the latent space from data (Bishop, 1999; Neal, 2012; Titsias and Lawrence, 2010). Similar to previous work in the field, we also use variational inference to facilitate arbitrary observation noise models, including non-Gaussian models more appropriate for electrophysiological recordings (Duncker and Sahani, 2018; Zhao and Park, 2017; Keeley et al., 2020b; Zhao et al., 2020; Liu and Lengyel, 2021; Schimel et al., 2021). While variational inference has proven a useful framework for such non-conjugate likelihood models, alternative approaches exist including the use of polynomial approximations to the non-linear terms in the likelihood (Keeley et al., 2020a). Another major challenge in the development of GP-based latent variable models such as bGPFA is to ensure scalability for longer time series. In this work, we utilize advances in variational inference (Kingma and Welling, 2014; Rezende et al., 2014) to facilitate scalability to the large datasets recorded in modern neuroscience. In particular, we contribute a new circulant variational GP posterior expressed partly in the Fourier domain that is both accurate and scalable. This is similar to Keeley et al. (2020b), who address the problem of scalability by assuming independence across Fourier features and formulating variational inference in the Fourier domain. However, we instead perform inference in the time domain and include additional factors in our variational posterior that ensure smoothness over time and allow for non-stationary posterior covariances. In contrast to these approaches, Zhao and Park (2017) rely on a low rank approximation to the prior covariance for inference and temporal subsamples for hyperparameter optimization to overcome the computational cost of model training. A conceptually similar approach employed by Duncker and Sahani (2018) is the use of inducing points which has been studied extensively in the Gaussian process literature (Hensman et al., 2015a, 2013; Titsias, 2009). However, such low rank approximations can perform poorly on long time series where the number of inducing points needed is proportional to the recording duration (Chang et al., 2020).

bGPFA is also closely related to Gaussian process latent variable models (GPLVMs) (Lawrence and Hyvärinen, 2005; Titsias and Lawrence, 2010) which have recently found use in the neuroscience literature as a way of modelling flexible, nonlinear tuning curves (Wu et al., 2017; Jensen et al., 2020; Liu and Lengyel, 2021). This is because integrating out the loading matrix ***C*** in *p*(***Y***|***X***) with a Gaussian prior gives rise to a Gaussian process with a linear kernel. The low-rank structure of this linear kernel yields computationally cheap likelihoods, and our variational approach to estimating log *p*(***Y***|***X***) is in fact equivalent to the sparse inducing point approximation used in the stochastic variational GP (SVGP) framework (Hensman et al., 2013, 2015a). In particular, our variational posterior is the same as that which would arise in SVGP with at least *D* inducing points irrespective of where those inducing points are placed (Appendix H). We also note that for a Gaussian noise model, the resulting low-rank Gaussian posterior is the form of the exact posterior distribution (Appendix G). Additionally, since the bGPFA observation model and prior over latents are both GPs, bGPFA is an example of a deep GP (Damianou and Lawrence, 2013) with two layers – the first with an RBF kernel and the second with a linear kernel. Finally, our parameterizations of the posteriors *q*(***x***_*d*_) and *q*(***f**_n_*) can be viewed as variants of the ‘whitening’ approach introduced by Hensman et al. (2015b) which both facilitates efficient computation of the KL terms in the ELBOs and also stabilizes training (Section 2.2).

### Conclusion

In summary, bGPFA is an extension of the popular GPFA model in neuroscience that allows for regularized, scalable inference and automatic determination of the latent dimensionality as well as the use of non-Gaussian noise models appropriate for neural recordings. Importantly, the hyperparameters of bGPFA are efficiently optimized based on the ELBO on training data, which alleviates the need for cross-validation or complicated algorithms otherwise used for hyperparameter optimization in overparameterized models (Jensen et al., 2020; Wu et al., 2017; Yu et al., 2009; Keshtkaran and Pandarinath, 2019; Keshtkaran et al., 2021; Gao et al., 2016). Our approach can also be extended in several ways to make it more useful to the neuroscience community. For example, replacing the spike count-based noise models with a point process model would provide higher temporal resolution (Duncker and Sahani, 2018), and facilitate inference of optimal temporal delays across neural populations (Lakshmanan et al., 2015). This will likely be useful as multi-region recordings become more prevalent in neuroscience (Keeley et al., 2020c). Additionally, by substituting the linear kernel in *p*(***Y***|***X***) for an RBF kernel in Euclidean space (Wu et al., 2017) or on a non-Euclidean manifold (Jensen et al., 2020), we can recover scalable versions of recent GPLVM-based tools for neural data analyses with automatic relevance determination.

## Acknowledgements

We are grateful to O’Doherty et al. (2017) for making their data publicly available and to Marine Schimel and David Liu for insightful discussions. We thank Marine Schimel, Yashar Ahmadian, Peter Stone, and Jonathan So for helpful comments on the manuscript. We thank David Liu for contributions to the codebase used for our analyses. K.T.J. was funded by a Gates Cambridge scholarship and J.T.S. by a Churchill scholarship.

## Appendix

### A Further analyses of preparatory dynamics in the primate reaching task

We performed analyses as in Figure 3f using the raw data (***Y***) and using factor analysis (FA) with 20 latent dimensions instead of using the bGPFA latent states. The raw data ***Y*** showed a high degree of similarity at target onset compared to movement onset, but little discernable structure as a function of reach direction at either point in time (Figure 5a-b).

**Figure 5:**
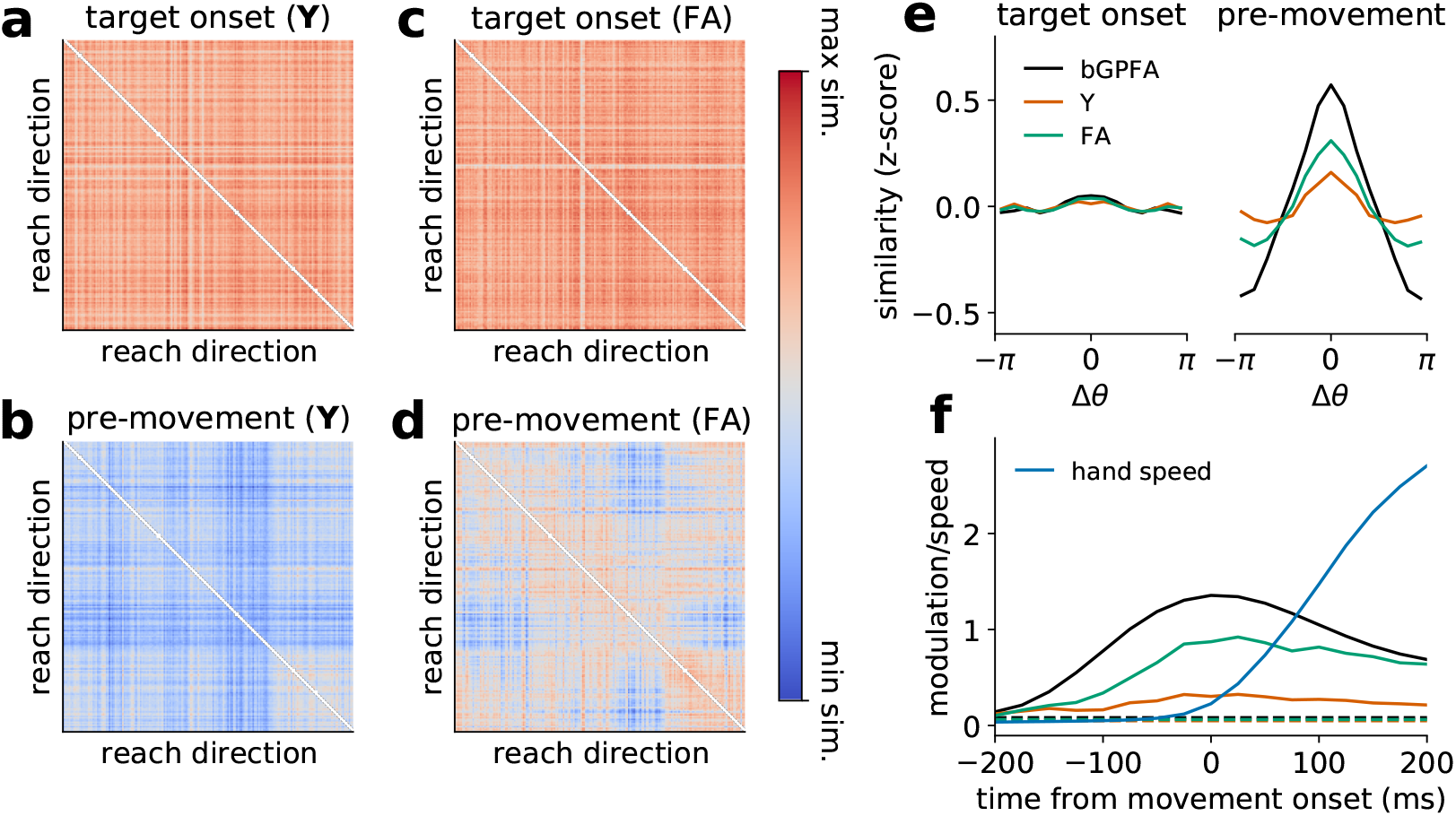
Further analyses of M1 preparatory dynamics. **(a-d)** Similarity matrix of raw neural activity ***Y*** (a & b) and latent states found by FA (c & d) at target onset (a & c) and 75 ms prior to movement onset (b & d), with analyses performed as in Figure 3f. **(e)** z-scored similarity as a function of difference in reach direction; here, the mean similarity across pairs of reaches is shown at target onset (left) and 75 ms prior to movement onset (right). The bGPFA latent states show much stronger modulation than either raw neural activity (***Y***) or latent states from FA. **(f)** Modulation of similarity by reach direction as a function of time from movement onset. Modulation was defined as the difference between maximum and minimum z-scored similarity as a function of difference in reach direction (peak-to-trough in panel e). Blue solid line indicates the z-scored hand speed, confirming the absence of premature movement relative to our definition of movement onset. bGPFA latent similarity increases well before hand speed and starts decreasing substantially before the hand speed peaks. Dashed lines indicate modulation at target onset for each method.

While the FA latent distances exhibited no modulation by reach direction at target onset, FA did discover weak modulation at movement onset (Figure 5a-b). This is qualitatively consistent with our results using bGPFA but with a lower signal to noise ratio. Here and in Section 3.2, we defined movement onset as the first time during a reach where the cursor velocity exceeded 0.025ms^−1^, and we observed little to no quantifiable movement before this point (Figure 5f). We also discarded ‘trials’ with premature movement for all analyses here and in Section 3.2, which we defined as reaches with a reaction time of 75 ms or less.

To quantify and compare how neural activity was modulated by the similarity of reach directions for different analysis methods, we first computed z-scores of the similarity matrices for both the bGPFA latent states, raw activity, and the latent states from FA. z-scores were calculated as *z* = (***S*** – *mean*(***S***))/*std*(***S***) for each similarity matrix ***S***, and the diagonal elements were excluded for this analysis. We then computed the mean of the z-scored pairwise similarities as a function of difference in reach direction across all pairs of 762 reaches. We found that none of the datasets exhibited notable modulation at target onset (Figure 5e). In contrast, the neural data exhibited modulation by reach similarity 75 ms prior to movement onset. This modulation was strongest for the bGPFA latent states followed by the FA latents, and the modulation by reach similarity was very weak for the raw neural activity (Figure 5e). To see how this modulation by reach direction varied as a function of time from movement onset, we computed the difference (*δz*) between the maximum and minimum of the modulation curves and repeated this analysis at different times prior to and during the reach process. We found that the modulation in neural activity space increased before any detectable movement, with bGPFA showing the strongest signal followed by factor analysis and then the raw activity (Figure 5f). Indeed, the bGPFA latent modulation was maximized near movement onset, while the reach speed did not peak until several hundred milliseconds after movement onset where bGPFA latent trajectories have started to diverge again. Taken together, these results confirm that our analyses of bGPFA preparatory states do not reflect premature movement onset, and that they are not artifacts of the temporal correlations introduced by our GP prior since noisier but qualitatively similar results arise from the use of factor analysis.

For further comparison with non-Bayesian Gaussian process factor analysis, we also fitted Poisson GPFA (P-GPFA) to the primate dataset using our variational inference approach for scalability but without a prior over ***C*** (Section 2.1). For this analysis, we used 16 latent dimensions as inferred by bGPFA, and we subselected latent processes with timescales ≤ 200 ms to study the putatively fast motor preparation as for bGPFA. We then orthogonalized the latent dimensions by performing an SVD on the loading matrix (see Yu et al., 2009 for details). Similar to our results from bGPFA, we found that the latent trajectories became modulated by movement direction prior to movement onset (Figure 6a) with a similar degree of modulation to bGPFA (*δz* = 1.01 for bGPFA; *δz* = 0.99 for P-GPFA). When visualizing the latent trajectories for the example reaches considered in Figure 3e, we also found that these diverged by reach direction (Figure 6b), similar to the bGPFA latent trajectories and unlike vanilla factor analysis which assumes temporal independence *a priori* (Figure 6c)

**Figure 6:**
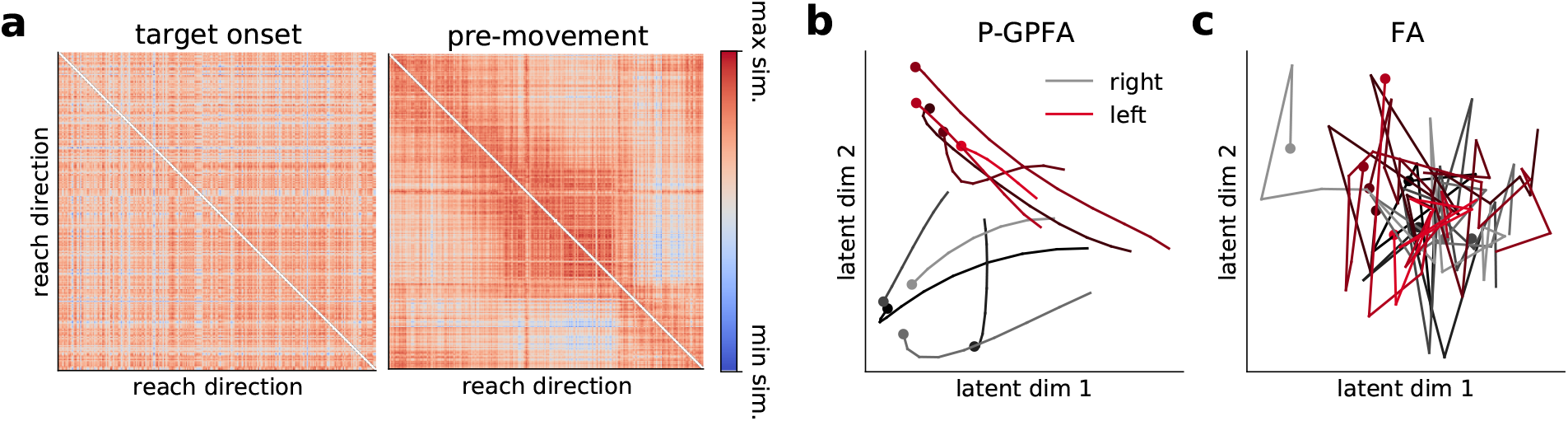
Analyses of M1 dynamics with GPFA and FA. **(a)** P-GPFA was fitted to data recorded from M1 during the self-paced reaching task. We computed the similarity matrix of the latent state at stimulus onset showing no obvious structure (left) and 75 ms prior to movement onset showing modulation by reach direction (right). Reaches are sorted by reach direction along both axes. **(b)** Example P-GPFA latent trajectories in the two principal latent dimensions for five rightward reaches (grey) and five leftward reaches (red). Trajectories are plotted from the appearance of the stimulus until movement onset (circles; the trajectories shown are the same as in Figure 3e). **(c)** As in (b), now showing latent trajectories inferred by factor analysis. These exhibit less discernable structure due to the lack of an explicit smoothness prior.

### B S1 activity

In this section, we compare the latent processes inferred for M1 dynamics to those inferred by applying bGPFA to the recordings from S1. In contrast to the clustering by reach direction in M1, there was less obvious modulation by movement direction in S1 prior to movement onset (Figure 7). To quantify this, we again computed the degree of modulation prior to movement onset, which was *δz* = 0.38 for S1 compared to *δz* = 1.01 for M1. This is also consistent with our decoding analyses in Figure 3b which showed that activity in M1 predicts movement 100-150 ms into the future while S1 activity is not predictive of future kinematics to the same extent.

**Figure 7:**
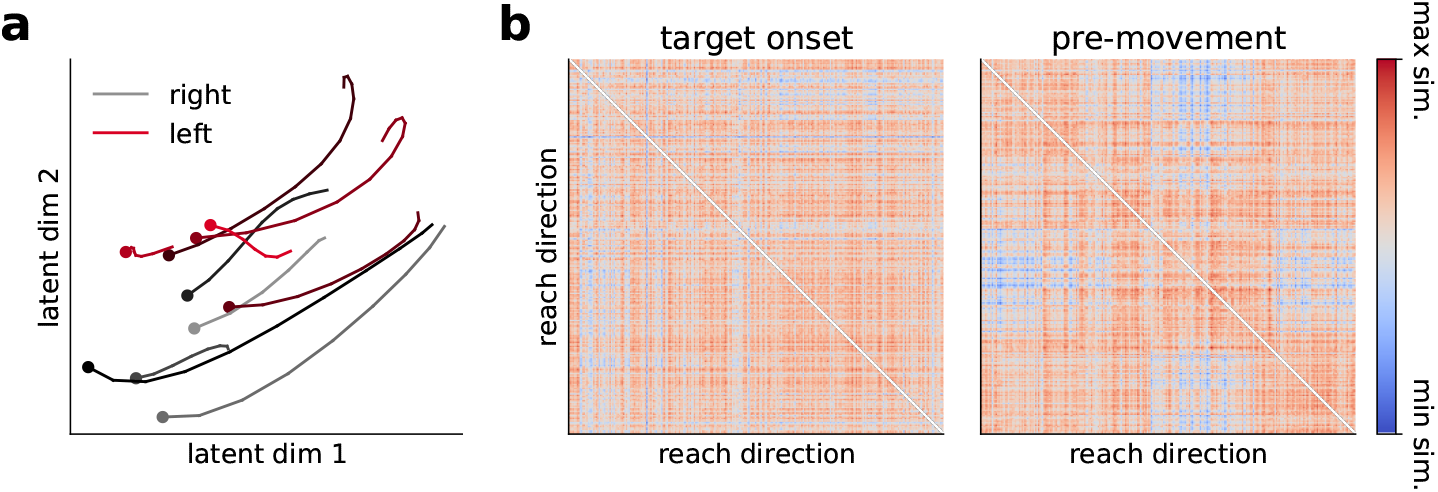
Analyses of S1 dynamics by reach direction. **(a)** bGPFA was fitted to data recorded from S1 during the self-paced reaching task. The panel shows example latent trajectories in the two most informative latent dimensions for five rightward reaches (grey) and five leftward reaches (red). Trajectories are plotted from the appearance of the stimulus until movement onset (circles; the trajectories shown are the same as in Figure 3e). Unlike the M1 latent trajectories, there is no obvious clustering by reach direction during movement preparation. **(b)** Similarity matrix of the latent state at stimulus onset (left) and 75 ms prior to movement onset (right). Reaches are sorted by reach direction along both axes, and there is no obvious structure in either similarity matrix in contrast to the results for the M1 recordings.

### C Further reaction time analyses

For analyses of correlations between latent distances and reaction times, we only considered reaches with a reaction time of at least 125 ms and at most 425 ms, which retained 712 of 762 reaches (Figure 8a). This is because very long reaction times may reflect the monkey not being fully engaged with the task during those reaches, and very short reaction times may reflect spurious movement. To confirm that our finding of a strong correlation between latent distance and reaction time in Figure 3g is not an artifact of the temporal correlations introduced by the bGPFA generative model, we generated a synthetic control distribution. Here we drew 50,000 synthetic latent trajectories from our learned generative model with trajectory durations matched to those observed experimentally on each trial. We then computed mean preparatory states and latent distances to preparatory states as in the experimental data (Section 3.2) and computed correlations with the experimental reaction times. We found a mean correlation of 0.02 and a range of −0.14 to 0.18 in the synthetic data, suggesting that our generative model may introduce weak correlations between latent distances and reaction times. However, the experimentally observed correlation of 0.45 was much larger than what could be expected by chance. This verifies our finding that the distance from the latent state at target onset to the corresponding preparatory state has behavioral relevance, with better initial states leading to shorter reaction times.

**Figure 8:**
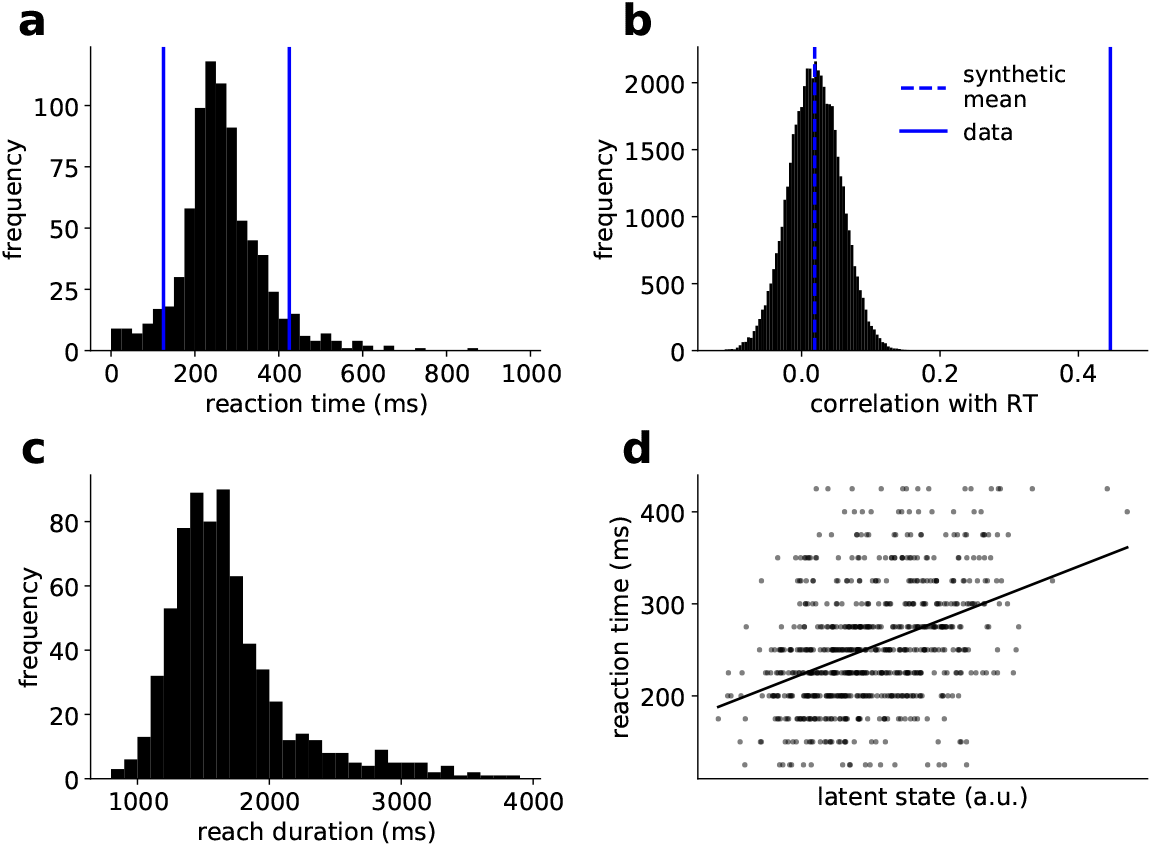
Further reaction time analyses. **(a)** Histogram of reaction times across all succesful reaches. For our correlation analyses, we only considered reaches with a reaction time between 125 ms and 425 ms (blue vertical lines). **(b)** Pearson correlations between distance to prep state and reaction time in synthetic data. Histogram corresponds to correlations between the true reaction times and 50,000 draws from the learned generative model. Blue dashed line indicates mean across all synthetic datasets (0.02), which is much smaller than the observed correlation in the experimental data of 0.45 (blue solid line). **(c)** Histogram of reach durations for all reaches with a reaction time between 125 ms and 425 ms. **(d)** Plot of reaction time against the value of a long timescale latent dimension at target onset (*τ* =1.4 s, *ρ* = 0.40).

Although we already find a fairly strong relationship between latent distances and reaction times, it is worth noting that several additional considerations may further improve such predictions. Notably, our naïve Euclidean distance metric could be improved by instead defining a metric based on the probabilistic model itself (Tosi et al., 2014). Additionally, while we categorize reaches by reach direction, reaches in the same direction can still have different start and end points on the grid (Figure 3a), leading to different posture and muscle activations which is likely to significantly affect neural activity. Our analysis by reach direction therefore only represents a coarse categorization of the rich behavioral space, and it remains to be seen how neural activity and latent trajectories are affected by e.g. posture during the task.

Finally we considered how the dynamics of long-timescale latent processes relate to the reaction time across trials (c.f. Section 3.3). Here we found that the two slowest dimensions had timescales of *τ* = 1.4 s and *τ* = 1.7 s, similar to the timescale of single reaches which generally lasted between 1 and 2 seconds (Figure 8c). Intriguingly, the latent state in these dimensions at target onset was predictive of reaction time with Pearson correlations of *ρ* = 0.40 and *ρ* = 0.36 respectively (Figure 8d). While the information about reaction time contained in these two dimensions was largely redundant, it was orthogonal to that encoded by the distance to preparatory state in the fast dimensions. In particular, a linear model had 19.9% variance explained from the distance to prep in fast dimensions, 15.7% variance explained from the slow latent dimension with the strongest correlation, and 28.7% when combining these two features which corresponds to 80.7% of the additive value.

### D Latent dimensionality

In this section, we estimate the dimensionality of the primate data as a function of the offset between M1 and S1 spike times using both bGPFA and participation ratios computed on the basis of PCA (Recanatesi et al., 2019). The participation ratio is defined as

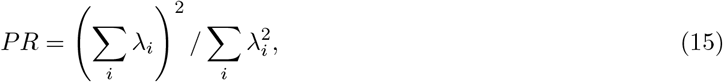

where λ*_i_* is the *i^th^* eigenvalue of the covariance matrix ***YY**^T^*. When computing the participation ratio of the data as a function of the M1 spike time shift, we find that the dimensionality is minimized for a shift of 75-100 ms (Figure 9). This suggests that the neural recordings can be explained more concisely when taking into account the offset in decoding between M1 and S1 which is consistent with the increased log likelihood after shifting the M1 spikes (Section 3.2).

**Figure 9:**
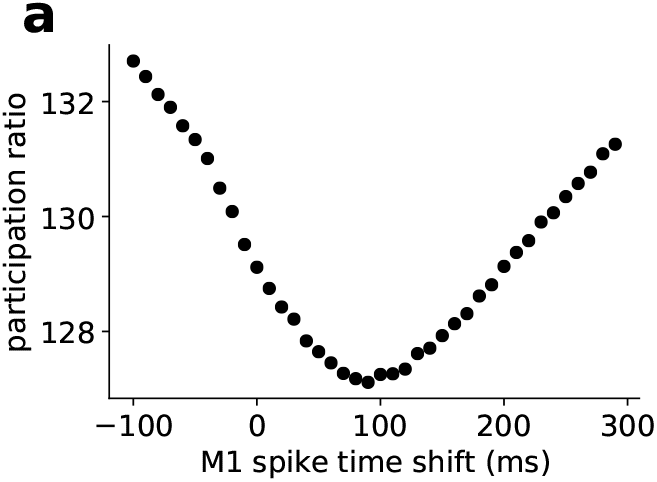
Neural dimensionality. **(a)** Participation ratio (Equation 15) as a function of temporal offset added to M1 spike times in the primate dataset.

**Figure 10:**
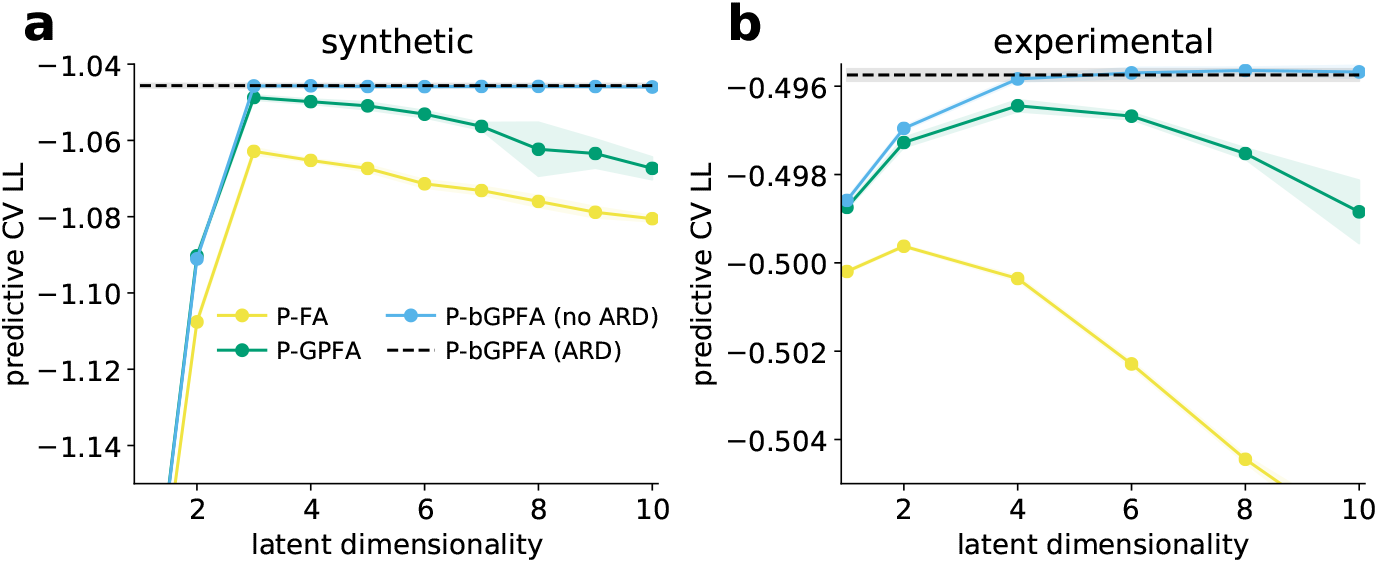
Bayesian GPFA applied to spike count data. **(a)** Cross-validated predictive log likelihoods of factor analysis (yellow), GPFA (green), and Bayesian GPFA without ARD (blue) fitted to synthetic data with a ground truth dimensionality of three for different model dimensionalities. All methods used a Poisson observation model *p*(*y_nt_*|*f_nt_*). bGPFA with ARD recovered the performance of the optimal non-ARD models without requiring a search over latent dimensionalities (black). **(b)** As in (a), now for models applied to a subset of the monkey reaching data analyzed in Section 3.2.

This trend is not directly observable in the number of dimensions retained by bGPFA with and without a 100 ms shift of the M1 spike times (18.8 ± 0.19 vs 18.7 ± 0.20 respectively across 10 model fits). However, the discrete nature of this dimensionality measure makes it relatively insensitive to small effects since it relies on stochastic differences in the retention of a dimension with little information content. We therefore utilized the interpretation of the learned prior scale 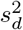 as a measure of variance explained (Appendix J) and defined a ‘participation ratio’ for bGPFA, similar to the PCA participation ratio considered above:

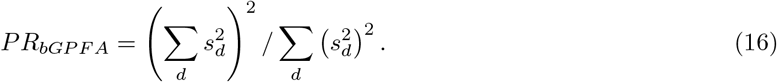

Here we found a strong effect of shifting the M1 spike times, which reduced the dimensionality from *PR_bGPFA_* = 6.16 ± 0.12 to *PR_bGPFA_* = 5.62 ± 0.06 (*p* = 0.001). Additionally, we note that bGPFA explains the data with only a handful of latent dimensions (19 total dimensions; 6 when re-weighted as a participation ratio). This is much lower than the dimensionality of 127-129 estimated by the PCA participation ratio which generally infers higher dimensionalities for noisier (more ‘spherical’) datasets.

### E Further validation of bGPFA on synthetic and biological data

In Figure 2a-b, we considered the performance of FA, GPFA and bGPFA on synthetic data with Gaussian noise. To further validate our method in a non-Gaussian setting relevant to the study of electrophysiological recordings, we also performed similar analyses on (i) synthetic data with Poisson noise and (ii) experimental recordings from the primate reaching task. In both cases, we compared FA, GPFA and bGPFA with Poisson noise, since such Poisson noise models are common in the neuroscience literature (Macke et al., 2012; Wu et al., 2017; Pandarinath et al., 2018; Zhao and Park, 2017). We note that these non-conjugate models are all readily implemented within our inference framework as special cases of bGPFA.

#### Synthetic data

In this section, we perform analyses similar to Figure 2b to validate bGPFA and its capacity for automatic relevance determination on synthetic data. We first generated a dataset drawn from the GPFA generative model but with a Poisson noise model after passing the activations through a softplus nonlinearity

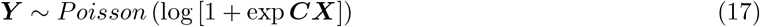

We then fitted Factor analysis, GPFA, and bGPFA and without ARD to the resulting dataset, all with Poisson observation models and exponential non-linearities (Appendix K; we denote these Poisson models as ‘P-FA’ etc.). To quantify performance, we computed the cross-validated predictive log likelihood 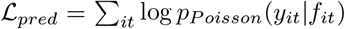, where *i* and *t* index neurons and time points in a held-out test set (results were similar when considering MSEs). When considering 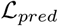 as a function of latent dimensionality, we found that both P-FA and P-GPFA exhibited a clear maximum at the true dimensionality of *D*\* = 3 while P-bGPFA without ARD was robust to overfitting, similar to our findings for the Gaussian models (Figure 2b). Finally, bGPFA with ARD was capable of automatically recovering this dimensionality as well as the maximum predictive performance achieved across the other models.

#### Experimental data

We then proceeded to perform an analysis as above on data from the self-paced monkey reaching dataset (O’Doherty et al., 2017). For this analysis, we used a smaller subset of the data for computational convenience, considering only 1000 timepoints but including all 200 neurons. We performed these analyses in 10-fold cross-validation, averaging performance over folds and repeating the entire analysis across 5 different random seeds. We again fitted P-FA and P-GPFA and found that these models exhibited a clear maximum in their predictive log likelihoods. As for the synthetic data, P-bGPFA without ARD was robust to overfitting, and the introduction of ARD allowed us to infer the optimal latent dimensionality of *D*\* ≈ 4 as well as achieving optimal performance without *a priori* assumptions about the latent dimensionality. The robustness to overfitting of bGPFA both with and without ARD suggests that it could also be a valuable tool in settings with large simultaneous recordings of thousands of neurons, which are becoming increasingly relevant with recent advances in neural recording technologies (Steinmetz et al., 2021; Pachitariu et al., 2017).

Taken together, these results further validate the utility of bGPFA on both synthetic and biological data with non-conjugate noise models. They also highlight the utility of automatic relevance determination in practice, where it obviates the need to perform extensive cross-validation to select an appropriate latent dimensionality for the experimental data.

### F Parameterizations of the approximate GP posterior

In this section, we compare different forms of the variational posterior *q*(***X***) discussed in Section 2.2. For factorizing likelihoods, the optimal posterior takes the form

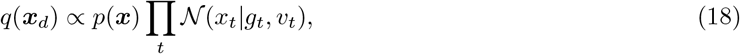

where *g_t_* and *v_t_* are variational parameters (Opper and Archambeau, 2009). Equation 18 might therefore seem to be an appropriate form of the variational distribution *q*(***X***). However, this formulation is computationally expensive and the likelihood *p*(***Y***|***X***) does not factorize across time in bGPFA.

Instead, we therefore consider approximate parameterizations of the form

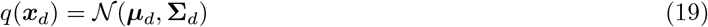

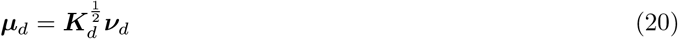

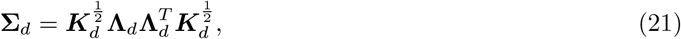

where 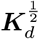 is a matrix square root of the prior covariance matrix ***K**_d_*, and 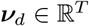 is a vector of variational parameters. This formulation simplifies the KL divergence term for each latent dimension in Equation 6 from

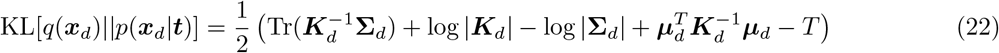

to

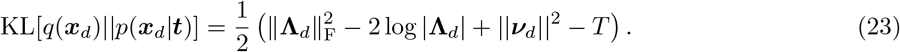

In the following, we drop the ·_*d*_ subscript to remove clutter, and we use the notation **Ψ** = diag(*ψ*_1_, …, *ψ_T_*) with positive elements *ψ_t_* > 0, to denote a positive definite diagonal matrix.

#### F.1 Square root of the prior covariance

For a stationary prior covariance ***K***, we can directly parameterize 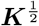 by taking the square root of *k*(·, ·) in the Fourier domain and computing the inverse Fourier transform. For the RBF kernel used in this work we get

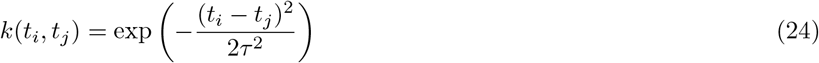

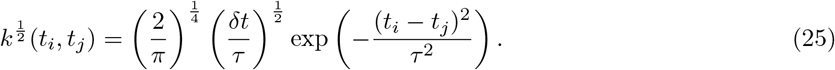

In this expression, *δt* is the time difference between consecutive data points, we have assumed a signal variance of 1 in the prior kernel, and we note that our parameterization only gives rise to the exact matrix square root of the RBF kernel in the limit where *T* ≫ *τ*. Note that this is the case in the present work since *T* ≈ 30 minutes is much larger than the longest timescales learned by bGPFA (*τ* ≈ 2 s). For most experiments in neuroscience, observations are binned such that time is on a regularly spaced grid and our parameterization can be applied directly. In other cases, kernel interpolation should first be used to construct a covariance matrix with Toeplitz structure (Wilson and Nickisch, 2015; Wilson et al., 2015).

#### F.2 Parameterization of the posterior covariance

We now proceed to describe the various parameterizations of **Λ** whose performance is compared in Figure 11. Other parameterizations are explored in Challis and Barber (2013).

**Figure 11:**
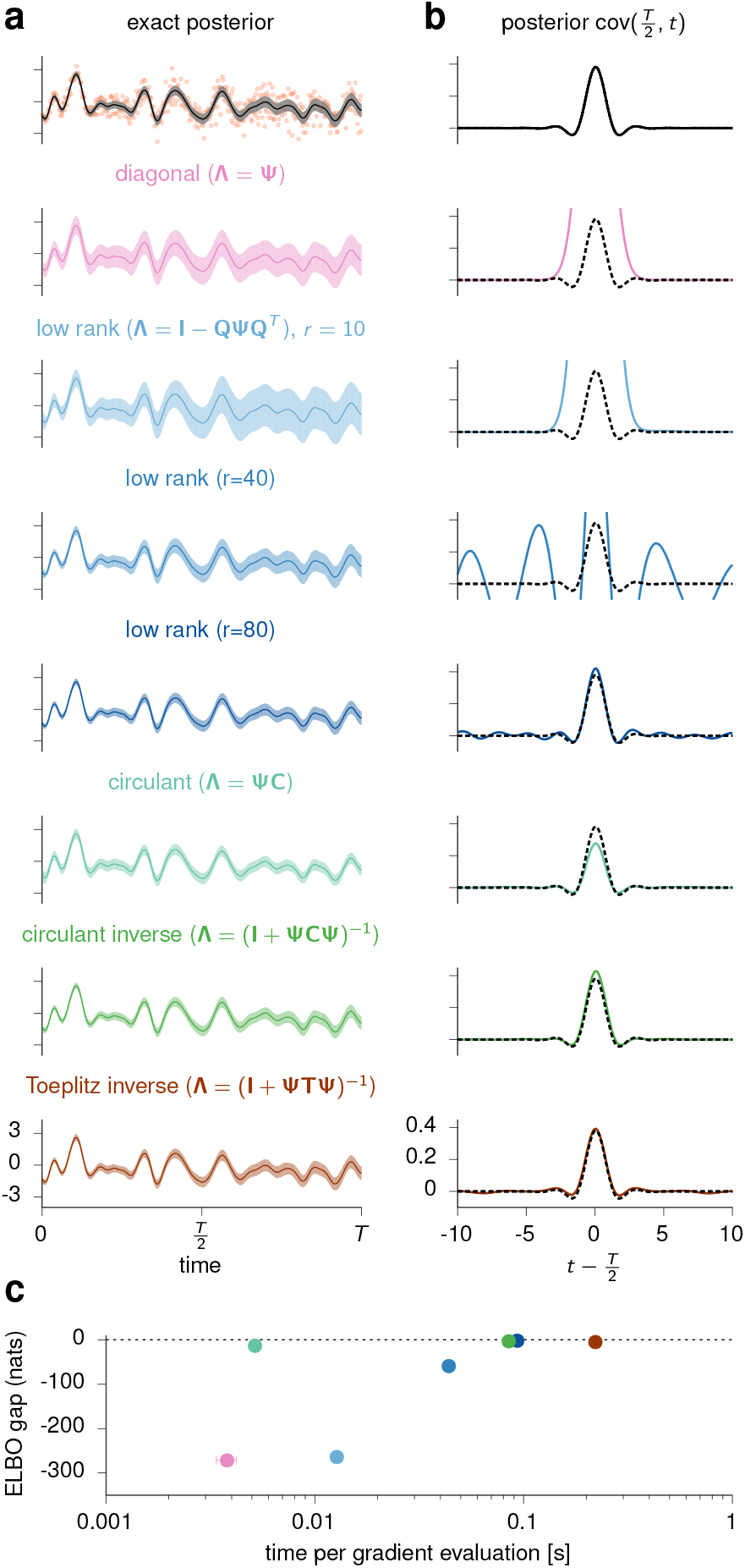
Comparisons of different forms of the approximate posterior *q*(*x*). **(a)** Synthetic data (orange dots) plotted together with the exact posterior (black) as well as the variational posteriors inferred by each whitened parameterization. The solid lines denote the (approximate) posterior means, and shaded areas indicate ±1 posterior standard deviations. **(b)** Slice through the posterior covariance (Cov_*x~q*(*x*)_ [*x*_*T*/2_, *x_t_*]) for the true posterior (top and black dotted lines) and the approximate methods. Each method has different characteristics, and the circulant parameterization provides a good qualitative fit at very low computational cost. **(c)** We defined the ‘ELBO gap’ of each method as ELBO — LL, where LL is the true data log likelihood. We plotted this against the time per gradient evaluation and found that the circulant parameterization achieved high accuracy with cheap gradients.

##### Diagonal Λ

We parameterize each latent dimension with **Λ** = **Ψ**. This gives rise to a KL term:

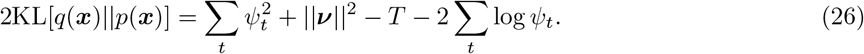

We can compute **Λ*v*** in linear time since **Λ** is diagonal which allows for cheap differentiable sampling:

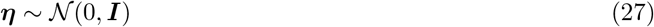

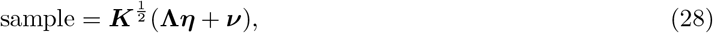

where the multiplication by 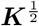 is done in 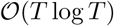 time in the Fourier domain.

##### Circulant Λ

We parameterize each latent dimension with **Λ** = **Ψ*C***. Here, 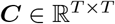 is a positive definite circulant matrix with 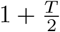 (integer division) free parameters, which we parameterize directly in the Fourier domain as 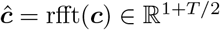, where ***c*** is the first column of ***C*** with 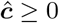 elementwise. We compute the KL as

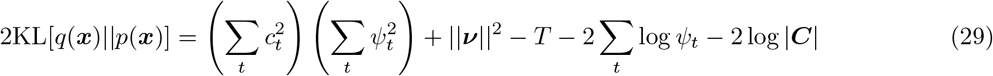

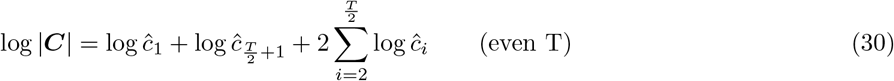

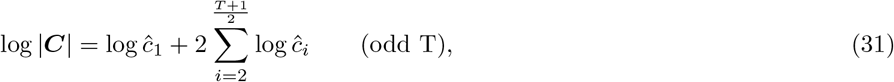

where 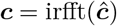. We can sample differentiably in 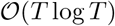 time by computing

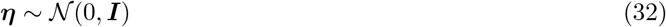

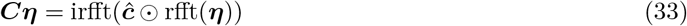

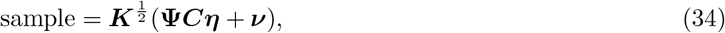

where ⊙ denotes the complex element-wise product.

##### Low-rank Λ

We let 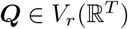 such that ***Q**^T^ **Q*** = ***I**_r_* and write

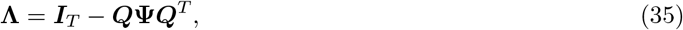

where we now constrain 0 < *ψ_i_* < 1 to maintain the positive definiteness of **Λ**. In practice, we keep **Q** on the Stiefel manifold (i.e. ***Q**^T^**Q*** = ***I**_r_*) by (differentiably) computing the QR decomposition of a *T* × *r* matrix of free parameters.

##### Circulant inverse Λ

We let ***C*** be a circulant positive definite matrix as above and parameterize

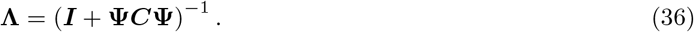

Computing **Λ*v*** products is done using the conjugate gradient algorithm, taking advantage of fast products with **Ψ** and ***C***; the same algorithm is also used to stochastically estimate log |**Λ**| and its gradient (see the appendix of Rutten et al., 2020).

##### Toeplitz inverse Λ

This proceeds just as for the circulant inverse form, with the circulant matrix ***C*** replaced by an arbitrary Toeplitz matrix (also exploiting fast ***Tv*** products):

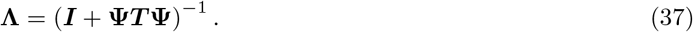

#### F.3 Numerical comparisons between different parameterizations

To compare these parameterizations, we generated a synthetic dataset (Figure 11a, orange dots) over *T* = 1000 time bins by drawing samples {*y*_1_, …, *y_T_*} as *y_t_* = *x_t_* + *σ_t_ξ_t_* where 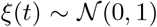 with non-stationary *σ_t_* growing linearly from 0.1 to 0.5 over the whole range 0 ≤ *t* < *T*, and 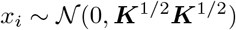 with ***K***^1/2^ given by Equation 25. We fixed these generative parameters to their ground truth and optimized the ELBO w.r.t. the variational parameters in this simple regression setting. We found that all of the parameterizations accurately recapitulated the GP posterior mean (Figure 11a). However, the degree to which they captured the non-stationary posterior covariance and data log likelihood varied between methods (Figure 11b-c). To quantify this, we computed the difference between the asymptotic ELBO of each method and the exact log marginal likelihood. This ELBO gap was small for the circulant parameterization, the inverse methods, and the low rank parameterization with sufficiently high *r*. Although the circulant parameterization did not fully capture the non-stationary aspect of the posterior variance, this did not affect the ELBO gap substantially. Importantly, however, the circulant parameterization was more than an order of magnitude faster per gradient evaluation than the other methods with comparable accuracy (Figure 11c). For these reasons as well as the excellent performance in a latent variable setting (Section 3.1, Section 3.2, Appendix E), we used the circulant parameterization for all experiments. However, repeating all analyses with a simple diagonal parameterization also lead to good performance and qualitatively similar results.

### G Relation between variational posterior over *F* and true posterior

Here we show that our parameterization of *q*(***f**_n_*) includes the exact posterior in the case of Gaussian noise. When the noise model is Gaussian (i.e., 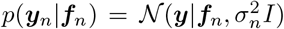), we can compute the posterior over 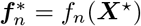 at locations ***X***^⋆^ in closed form:

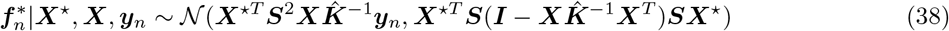

where 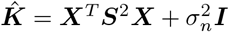. Note that the posterior is low-rank as the rank of 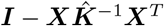 is at most *D*. This means that when we do variational inference, we can parameterize our approximate posterior as:

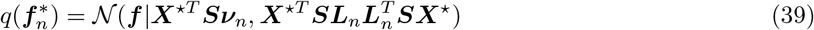

where 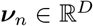 and 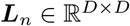 are the parameters of the approximate posterior (Section 2.2). We see that this parameterization is exact when:

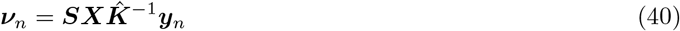

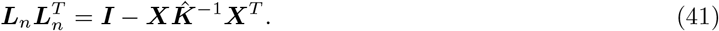

Note that the right-hand side of Equation 41 is guaranteed to be positive definite because the true posterior must be positive definite. Importantly, for this parameterization, the KL term in Equation 10 simplifies to

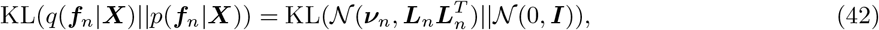

which is independent of ***X*** and allows us to do efficient inference due to the low dimensionality of ***ν**_n_* and ***L**_n_*.

### H Relation between variational posterior over *F* and SVGP

For general non-Gaussian noise models, the parameterization in Appendix G will no longer be exact. However, here we show that it is in this case equivalent to a stochastic variational Gaussian process (SVGP; Hensman et al., 2013). In SVGP, we choose a variational distribution:

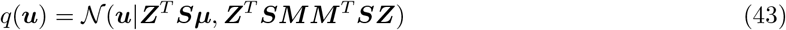

at inducing points 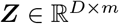, where ***μ*** and ***M*** are the “whitened” parameters (Hensman et al., 2015). This gives an approximate posterior:

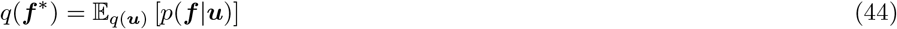

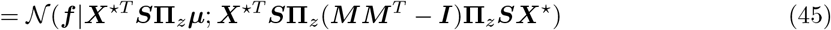

where **Π***_z_* = ***SZ***(***Z**^T^**S***^2^***Z***)^−1^***Z**^T^**S***. If we choose *m* = *D* inducing points such that 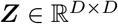 and make sure ***Z*** has full rank, then ***Π**_z_* = ***I*** and thus

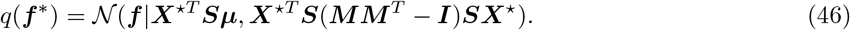

We recover the parameterization in Section 2.2 when

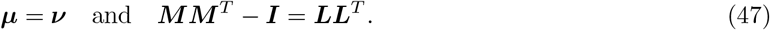

For these more general noise models, the whitened parameterization of *q*(***f***) still gives rise to a computationally cheap KL divergence that is independent of ***X*** as in Equation 42:

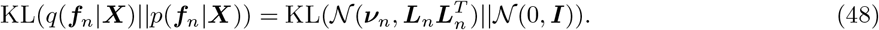

In summary, we have shown that (i) our parameterization of *q*(***f**_n_*) has sufficient flexibility to learn the true posterior when the noise model is Gaussian (Appendix G), and (ii) it is equivalent to performing SVGP where the locations of the inducing points do not matter provided that their rank is at least as high as the number of latent dimensions.

### I Automatic relevance determination

Here we briefly consider why introducing a prior over the factor matrix enables automatic relevance determination. These ideas reflect results by Bishop (1999) and our experiments in Section 3.1.

For simplicity, we will first consider the case of factor analysis where 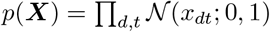. This gives rise to a marginal likelihood (with Gaussian noise) equal to

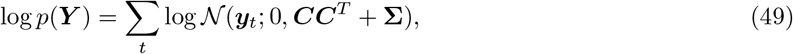

where 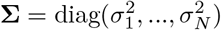 is a diagonal matrix of noise parameters. It is in this case quite clear that the optimal marginal likelihood is a monotonically increasing function of the latent dimensionality, since any marginal likelihood reachable with a certain rank *D* is also reachable with a larger rank *D*′ > *D*; increasing *D* can only increase model flexibility. We could in this case threshold the magnitude of the columns of ***C*** to subselect more ‘informative’ dimensions, but this is not inherently different from putting an arbitrary cut-off on the variance explained in PCA, and there is no Bayesian “Occam’s razor” built into the method (MacKay, 2003).

Consider now the case where we put a unit Gaussian prior on *c_nd_*. In this case, {*c_nd_*} are no longer parameters of the model but rather latent variables to be inferred, which intuitively should reduce the risk of overfitting. To expand on this intuition, consider the ELBO (c.f. Section 2.1) that results from introducing such a prior over *c_nd_*:

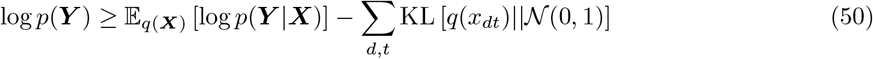

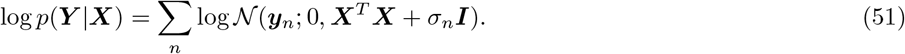

Here we see that if a dimension *d* is truly uninformative, it should have *x_dt_* = 0 ∀*_t_* to avoid contributing noise to the likelihood term via ***X**^T^**X***. However, reducing this noise will increase the prior KL term, driving it to infinity in the limit of zero noise since the variational posterior over the *d^th^* latent at time *t*, *q*(*x_dt_*), is in this case a delta function at zero. Optimizing the ELBO therefore involves a balance between mitigating the noise induced by ***X**^T^**X*** and reducing the KL penalty, with both of these terms contributing to a decreased ELBO compared to the model without uninformative dimensions. Thus the prior over *c_nd_* counteracts the overfitting that would normally occur when increasing the latent dimensionality in classical factor analysis, and this Bayesian treatment will lead to a decrease in the ELBO with increasing dimensionality beyond the optimal *D*^⋆^ that is needed to adequately explain the data.

Finally let us consider the case where we learn the prior scale of the factor matrix, such that 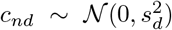 with *s_d_* optimized w.r.t. the ELBO. Critically, the likelihood term now becomes:

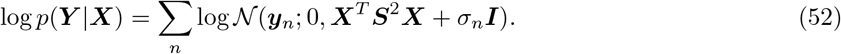

with ***S*** = diag(*s*_1_, …, *s_D_*). In this case, adding uninformative dimensions beyond the optimal *D*^⋆^ still cannot increase the ELBO (in the limit of large *N*). However, letting *s_d_* → 0 for these superfluous dimensions will prevent them from contributing to *p*(***Y***|***X***), thus allowing 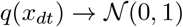 to drive the prior KL term to zero for these dimensions. In this limit, we recover both the ELBO and the posteriors associated with the *D*^⋆^-dimensional model. We thus have a built-in Occam’s razor which will shave off any uninformative latent dimensions, and these will be identifiable as dimensions for which *s_d_* ≈ 0 and 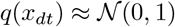.

These ideas generalize to GPFA where the posterior over latents will instead approach the GP prior 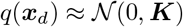 for uninformative dimensions. This corresponds to the limit of ***v*** → **0**, ***C*** → ***I***, and **Ψ** → ***I*** in our circulant parameterization in Section 2.2 and Appendix F. In all of our simulations, we found a clear clustering of dimensions after training with some clustered near zero *s_d_*, and others clustered with much larger *s_d_* (Figure 2c and Figure 3b). Note that in practice we do not actively truncate the model by discarding dimensions with *sd* ≈ 0 but merely use the terminology to indicate that these dimensions have negligible contributions to the posterior predictive *q*(***y***_*n*_), as well as to the latent posteriors *q*(***x**_d_*) for the dimensions with large *s_d_*.

### J Most informative dimensions

In this work, we refer to the latent dimensions with the highest values of *s_d_* as the ‘most informative dimensions’. We do this because (i) observing the value of the corresponding latent *x_d_* decreases the variance of the expected distribution of neural activity more as *s_d_* increases, and (ii) the Fisher information of *x_d_* increases as *s_d_* increases.

To show this, we consider how the distribution over *f_n_* (the activity of neuron *n*) given ***c**_n_* (the *n*^th^ row of ***C***) changes when *x_d_* (the value of the *d^th^* latent) is known, and how this varies with *s_d_*. In the following, we omit the ·_*n*_ subscript for notational simplicity, and we note that *f*, *x_d_* and *c_d_* are all scalar values. With unknown *x_d_*, *f* is Gaussian with zero mean and variance 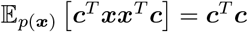. Thus,

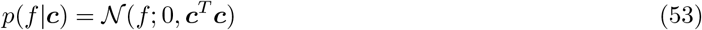

In contrast, for known *x_d_*, we have

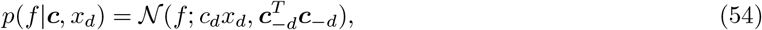

where ***c***_−*d*_ is ***c*** with the *d*^th^ element removed. We thus see that the decrease in variance of *f* from observing *x_d_* is 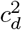. Finally, we can approximate the process of averaging this quantity over neurons by noting that 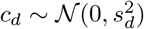 and marginalising out ***c***:

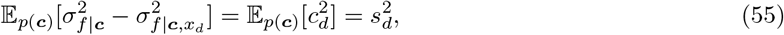

where 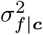 is the variance of *p*(*f*|***c***). Thus, 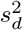 can be interpreted as the expected decrease in the variance of the denoised neural activity *f* when learning the value of the *d^th^* latent.

This can also be understood in information-theoretic terms by considering the Fisher information of the *d^th^* latent dimension which is given by

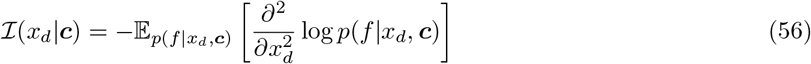

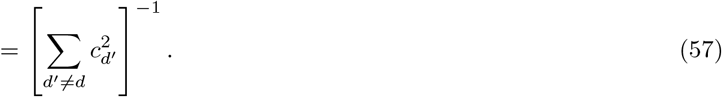

To relate this quantity to our prior scale parameters {*s_d_*}, we consider the expectation of the inverse Fisher information:

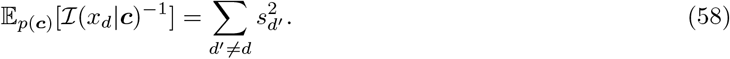

For a given set of latent dimensions [1, *D*] with corresponding 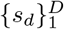, we thus see that the expected *inverse* Fisher information is *minimized* for the dimension with the highest value of *s_d_*. In Figure 2 and Figure 3 we use *s_d_* together with the posterior latent mean parameters ***v**_d_* to identify ‘discarded’ dimensions.

### K Noise models and evaluation of their expectations

#### Gaussian

The Gaussian noise model is given by

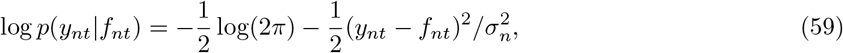

where *σ_n_* is a learnable parameter. In this case we can easily compute the expected log-density under the approximate posterior analytically:

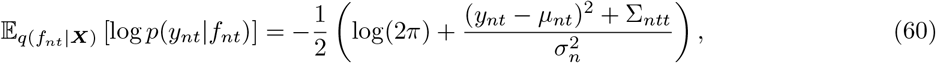

where 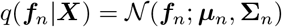 and Σ_*ntt*_ is the approximate posterior variance of neuron *n* at time *t* (i.e., the *t*^th^ diagonal element of **∑**_*n*_).

#### Poisson

The Poisson noise model is given by

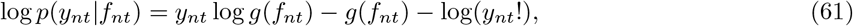

where *g* is a link function. If we choose an exponential link function (i.e., *g*(*x*) = exp(*x*)), we can compute in closed-form the expected log-density of the approximate posterior as:

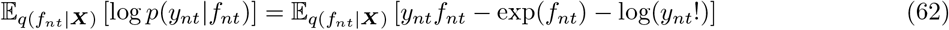

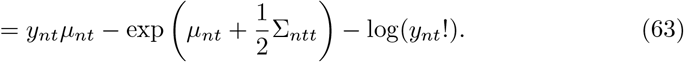

For the analyses shown in Figure 2c-d, we use the exponential link function.

For general link functions *g*, we may not be able to evaluate the expected log-density in closed-form. In this case, we approximate it with Gauss-Hermite quadrature:

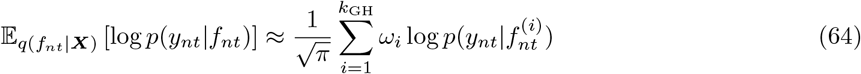

where

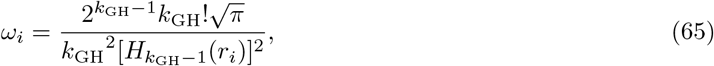

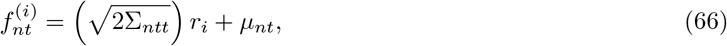

*H_k_*(*r*) are the physicist’s Hermite polynomials, and *r_i_* with *i* = 1, …, *k* are roots of *H_k_*(*r*). For a given order of approximation *k*_GH_, we can evaluate both *ω_i_* and *r_i_* using standard numerical software packages such as Numpy. In practice, we find that *k*_GH_ = 20 gives an accurate approximation to the expected log-density under the approximate posterior. Note that we could also estimate the expectation over *q*(*f_nt_*) for general link functions *g* using a Monte Carlo estimate, but we use Gauss-Hermite quadrature in this work since it has a lower computational cost and lower variance.

#### Negative binomial

The negative binomial noise model is given by

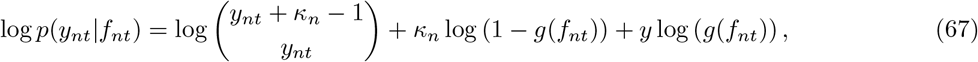

where *g*(*f_nt_*) denotes the probability of success in a Bernoulli trial. Here, each success corresponds to the emission of one spike in bin *t*, and thus *p*(*y_nt_*|*f_nt_*) is the distribution over the number of successful trials (spikes) before reaching *κ_n_* failed trials. The link function 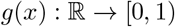 maps *f_nt_* to a real number between 0 and 1. In practice we use a sigmoid link-function *g*(*x*) = 1/(1 + exp(–*x*)).

In this model, *κ_n_* is a learnable parameter which effectively modulates the overdispersion of the distribution since the mean and variance of *p*(*y_nt_*|*f_nt_*) are given by:

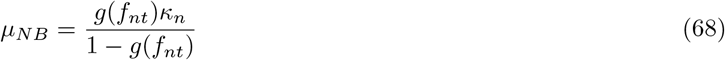

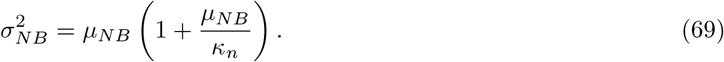

This is the parameter which we compare between the ground truth and trained models in Figure 2, and we see that the Poisson model is recovered for neuron *n* as *κ_n_* → ∞.

For the negative binomial noise model we cannot compute the expected log-density in closed-form. We instead approximate this expectation using Gauss-Hermite quadrature as described above.

### L Implementation

In this section, we provide pseudocode for bGPFA (Algorithm 1) with the circulant parameterization for *q*(***X***) and discuss other implementation details.

#### Algorithm 1: Bayesian GPFA with automatic relevance determination

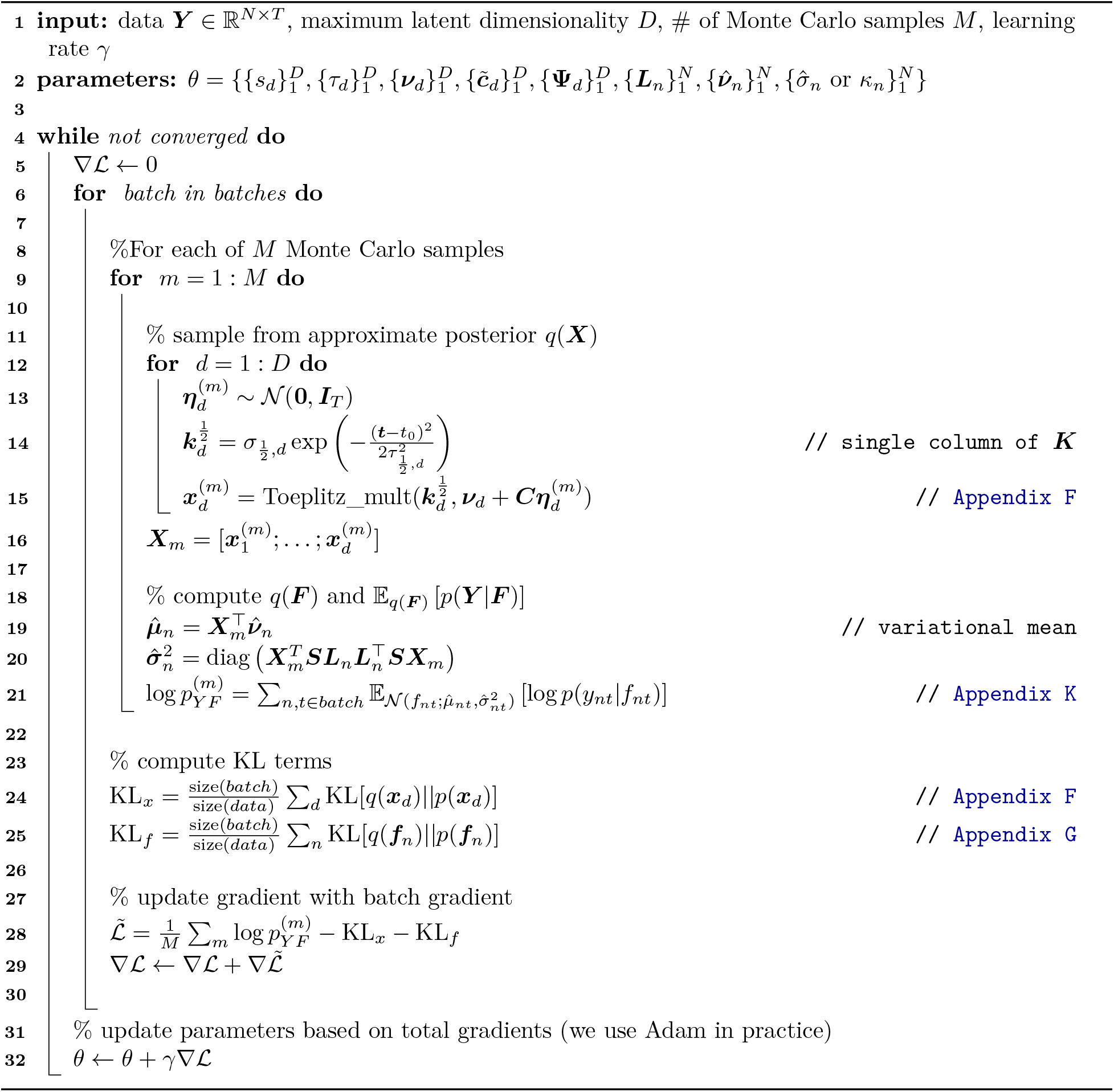

Note that we need to sample the full trajectory ***x**_d_* before subsampling for each batch due to the correlations introduced by ***K***. In practice, we run the optimization for 2000 passes over the full data which we found empirically lead to convergence of the ELBO. We used *M* = 20 Monte Carlo samples for each update step when fitting synthetic data and *M* =10 for the primate data. For all models, *q*(***X***) was initialized at the prior *p*(***X***). The prior scale parameters were initialized as 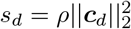 where ***c**_d_* is the *d^th^* row of the factor matrix ***C*** found by factor analysis (Pedregosa et al., 2011), and *ρ* = 3 was found empirically to give good convergence on the primate data. When using a Gaussian noise model, noise variances were initialized as the 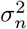 found by factor analysis. For negative binomial noise models, we initialized 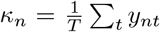 which matches the mean of the distribution to the data for *f* = 0. Length scales *τ* were initialized at 200 ms for all latent dimensions for the primate data and at ≈ 80% of the ground truth value for the synthetic data. Synthetic data was fitted on a single GPU with 8GB RAM. Primate data was fitted on a single GPU with 12GB RAM and took approximately 30 hours for a single model fit to the full dataset at 25 ms resolution. We also note that when fitting data with a Gaussian noise model, we mean-subtracted the original data, whereas we include explicit mean parameters in the Poisson and negative binomial noise models since they are non-linear (c.f. Appendix K).

#### Code availability

A PyTorch implementation of bGPFA is provided on [GitHub](https://github.com/tachukao/mgplvm-pytorch/tree/bgpfa).

### M Cross-validation and kinematic decoding

In this section, we describe the procedure for computing cross-validated errors in Figure 2, and performing kinematic decoding analyses in Figure 3. In these analyses, expectations over ***X*** were computed using the posterior mean of *q*(***X***) and expectations over ***F*** were computed using Monte Carlo samples from *q*(***F***).

#### Prediction errors

To compute cross-validated errors, we divide the time points into a training and a test set, 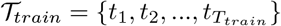 and 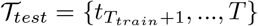, and similarly for the neurons 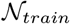 and 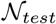. We also define 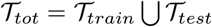 and 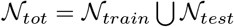. We first fit the generative parameters *θ_gen_* of each model to data from all the neurons at the training time points using variational inference (taking *θ* to include the parameters *ϕ* of *q_ϕ_*(***F***)):

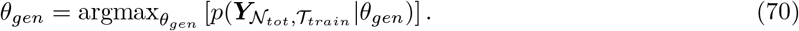

We then fix the generative parameters and infer a distribution over latents from the training neurons recorded at all time points using a second pass of variational inference:

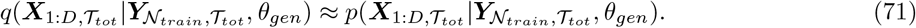

Finally we use the inferred latent states and generative parameters to predict the activity of the test neurons at the test time points

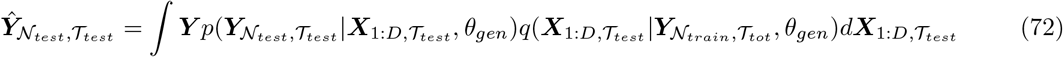

This allows us to compute a cross-validated predictive mean squared error as

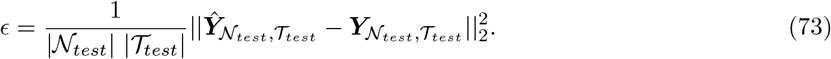

#### Kinematic decoding

For kinematic decoding analyses, we only considered the latents and behavior prior to a period of approximately 5 minutes where the monkey disengaged from the task (the first 1430 seconds; Section 3.3). Cursor positions in the x and y directions were first fitted with cubic splines and velocities extracted as the first derivative of these splines. To evaluate kinematic decoding performance, we followed Keshtkaran et al. (2021) and computed the expected activity of all neurons at all time points under our model:

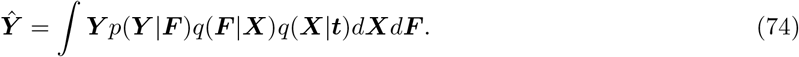

For non-Gaussian noise models, this can be viewed as the first non-linear step of a decoding model from the latent states ***X***. We then performed 10-fold cross-validation where 90% of the data was used to fit a ridge regression model which was tested on the held-out 10% of the data. The regularization strength was determined using 10-fold cross-validation on the 90% training data. The predictive performance was computed as the mean across the 10 folds. Models were fitted and evaluated independently for the hand x and y velocities, and the final performance was computed as the mean variance accounted for across these two dimensions. Results in Section 3.2 are reported as mean ± standard error across 10 different splits of the data into folds used for cross-validation.

## Notes

### Competing Interest Statement

The authors have declared no competing interest.

### Summary of Updates

NeurIPS 2021 camera-ready version.

